# Conserved prolines in the coiled coil-OB domain linkers of proteasomal ATPases facilitate eukaryotic proteasome base assembly

**DOI:** 10.1101/2020.11.13.381962

**Authors:** Chin Leng Cheng, Michael K Wong, Yanjie Li, Mark Hochstrasser

## Abstract

The proteasome is a large protease complex that degrades both misfolded and regulatory proteins. In eukaryotes, the 26S proteasome contains six different AAA+ ATPase subunits, Rpt1-Rpt6, which form a hexameric ring as part of the base subcomplex that drives unfolding and translocation of substrates into the proteasome core. Archaeal proteasomes contain only a single type of ATPase subunit, the proteasome-activating nucleotidase (PAN), which forms a trimer-of-dimers and is homologous to the eukaryotic Rpt subunits. A key PAN proline residue (P91) forms *cis* and *trans* peptide bonds in successive subunits around the ring, allowing efficient dimerization through upstream coiled coils. The importance of the equivalent Rpt prolines in eukaryotic proteasome assembly was unknown. We show an equivalent proline is strictly conserved in Rpt3 (in *S. cerevisiae*, P93) and Rpt5 (P76), well conserved in Rpt2 (P103), and loosely conserved in Rpt1 (P96) in deeply divergent eukaryotes, but in no case is its mutation strongly deleterious to yeast growth. However, the *rpt2-P103A*, *rpt3-P93A*, and *rpt5-P76A* mutations all cause synthetic defects with specific base assembly chaperone deletions. The Rpt5-P76A mutation decreases the levels of the protein and induces a mild proteasome assembly defect. The yeast *rpt2-P103A rpt5-P76A* double mutant has strong growth defects attributable to defects in proteasome base formation. Several Rpt subunits in this mutant form aggregates that are cleared, at least in part, by the Hsp42-mediated protein quality control (PQC) machinery. We propose that the conserved Rpt linker prolines promote efficient 26S proteasome base assembly by facilitating specific ATPase heterodimerization.

## Introduction

The eukaryotic 26S proteasome is a complex and highly abundant intracellular protease that comprises at least 33 different subunits; it uses the energy of ATP cleavage to unfold polyubiquitin-modified proteins and translocate them to a central chamber for proteolysis (1,2). The proteasome is composed of a 20S core particle (CP), which forms a barrel structure with a proteolytic chamber at its center, and a 19S regulatory particle (RP) on one or both ends of the CP. The RP is made up of two major sub-complexes, the lid and base, which can assemble independently. The lid contains a deubiquitylase subunit, Rpn11, that removes ubiquitin chains from substrates prior to their degradation. The base includes six distinct AAA+ ATPases (called Rpt1-6 in *Saccharomyces cerevisiae*) that form a heterohexameric Rpt ring in the order Rpt1-2-6-3-4-5 (2–4). The base has three additional non-ATPase subunits: Rpn1, Rpn2, and Rpn13 (Tomko and Hochstrasser 2013, Budenholzer et al. 2017).

Proteasome assembly must be carefully orchestrated due to the size, complexity, and abundance of this ~2.5 MDa complex. In eukaryotes, assembly of the base is facilitated by at least four dedicated chaperones: Nas2 (p27 in human), Nas6 (p28/gankyrin in human), Rpn14 (PAAF1 in human), and Hsm3 (S5b in human) (5–9). During base assembly, biochemical data suggest the Rpt subunits associate to form specific heterodimers along with their cognate chaperones: Hsm3-Rpt1-Rpt2 (and Rpn1), Nas2-Rpt4-Rpt5, and Nas6-Rpt3-Rpt6-Rpn14. These three “modules” then assemble into the ATPase ring. Adc17, an additional base assembly chaperone found only in yeast, is thought to bind directly to Rpt6 and facilitate Rpt3-Rpt6 dimerization, particularly under stress conditions when increased amounts of proteasomes are required (10). Expression of all proteasome base chaperones is also induced upon proteotoxic stress to enhance proteasome biogenesis (11).

In archaea, by contrast, ATPase ring assembly likely proceeds independently of dedicated chaperones. Instead of six paralogous ATPase subunits, the archaeal ATPase ring comprises six copies of a single AAA+ ATPase ortholog called the proteasome-activating nucleotidase (PAN) (12,13). The domain organization of PAN and the Rpts is conserved, beginning with an N-terminal coiled-coil (CC) domain followed by an oligonucleotide/oligosaccharide-binding (OB)-fold and the large and small domains typical of AAA+ ATPases (Figure 1A) (14,15). Similar to Rpt1, Rpt2, Rpt3 and Rpt5, PAN also contains a C-terminal HbYX (hydrophobic-Tyr-any residue) motif that engages surface pockets between the αsubunits of the outer heptameric rings of the CP (16,17).

**Figure 1.**
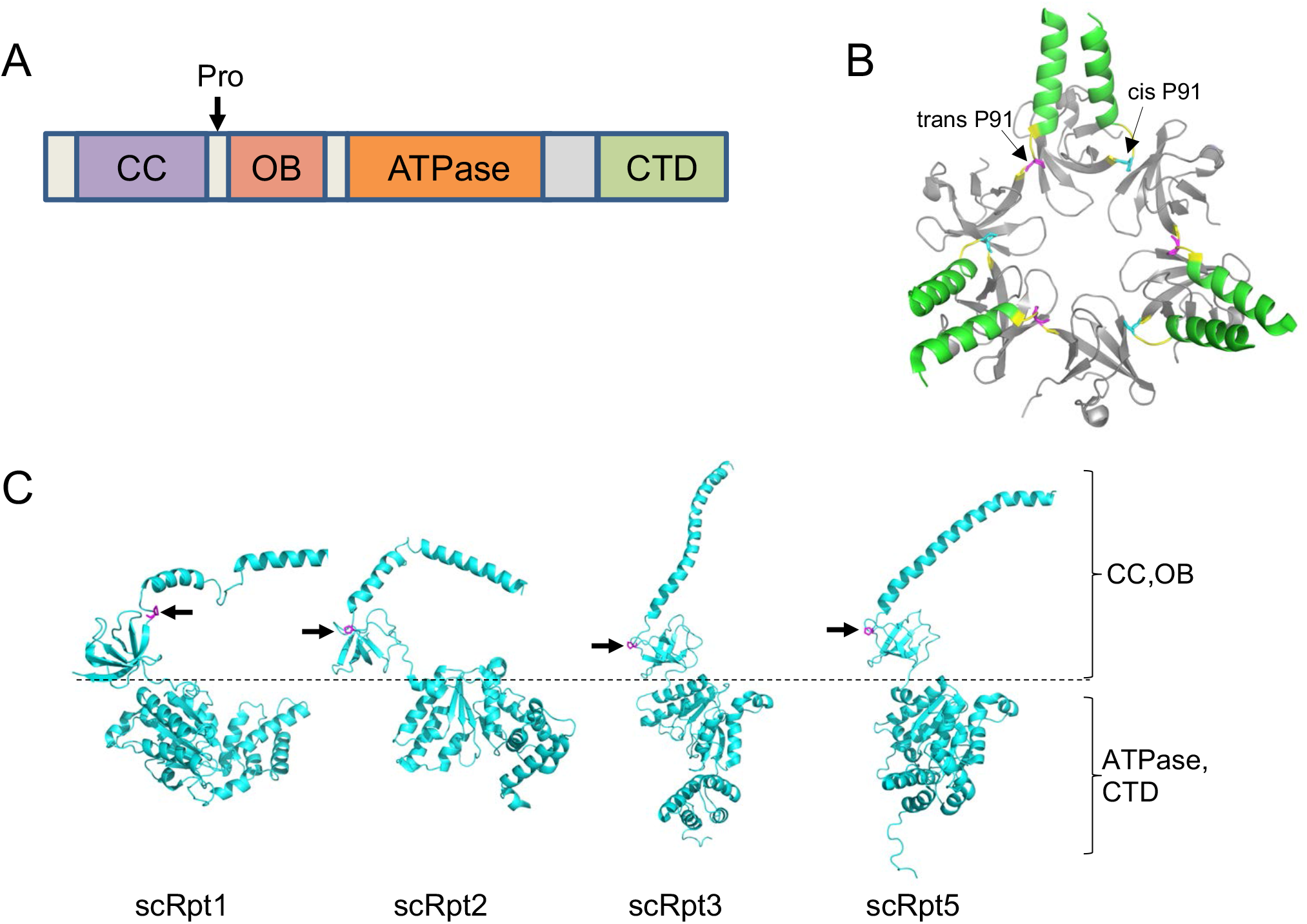
Structures of *Methanocaldococcus jannaschii* PAN (CC and OB domains) and *S. cerevisiae* Rpt1, Rpt2, Rpt3, and Rpt5, highlighting position of conserved proline residue. (A) Domain organization of PAN/Rpt subunit. Position of proline (if present) is indicated. CC, coiled coil; OB, oligonucleotide/oligosaccharide-binding domain; CTD, C-terminal domain characteristic for AAA+ ATPases. (B) Structure of *M. jannaschii* PAN (PDB ID:3H43). *trans* Pro91 residue is highlighted in magenta and *cis* Pro91 in blue. (C) *S. cerevisiae* Rpt subunits with conserved linker prolines from cryo-EM structure (PDB:5MP9). Linker prolines indicated in magenta.

In *Methanocaldococcus jannaschii*, the PAN ring arranges in a trimer-of-dimers configuration (14). Crystal structures of the N-terminal CC-OB segment of PAN revealed that the formation of dimers is dictated by the ability of the peptide bond preceding a specific proline residue, P91, in the short linker between CC and OB domains to adopt a *cis* conformation in one subunit of the dimer and *trans* conformation in the other (Figure 1B) (14). Analysis of peptide bonds in available protein structures have revealed that 6.5% of total imide bonds (X-Pro peptide bonds) have a *cis* conformation while only 0.05% of all amide bonds (X-nonPro peptide bonds) are in a *cis* conformation (18). The higher abundance of *cis* isomers of proline is due to the lower energy difference between *cis* and *trans* isomers relative to other amino acids (19). Despite this small energy difference, interconversion between *cis* and *trans* conformations of proline is a slow process and can be rate-limiting for protein folding and unfolding (20,21). Cells encode multiple prolyl isomerases that catalyze this interconversion (22).

An attempt to characterize recombinant *M. jannaschii* PAN-P91A and PAN-P91G mutant proteins *in vitro* was unsuccessful because the complexes were unstable, further highlighting the importance of this residue in PAN ring assembly (14). One study investigated ATP-independent chaperone activity of a PAN homolog, called ARC, in the actinobacterial species *Rhodococcus erythropolis* and found that mutation of the conserved proline (P62) in ARC-N (consisting of CC and OB domains) significantly reduced the ability of the complex to inhibit aggregation of denatured citrate synthase and luciferase, suggesting that the conserved proline is important for activity (23).

The equivalent proline residue is found in Rpt1 (P96), Rpt2 (P103), Rpt3 (P93), and Rpt5 (P76) in *S. cerevisiae* (Figure 1C) (24). Despite the importance of this residue in archaeal and actinobacterial proteasomal ATPases, its significance in eukaryotic Rpt subunits remains unexplored. Based on the order of the Rpt subunits in the heterohexamer and their pairwise interaction during base assembly, Rpt2, Rpt3, and Rpt5 have been predicted to have their linker prolines in the *cis* conformation (4). High-resolution structures of 26S proteasomes using cryogenic electron microscopy (cryo-EM) have allowed visualization of subunit interactions within the proteasome and different conformational states (1). However, there is currently no consensus on the *cis-trans* configuration at these Rpt prolines based on available cryo-EM structures of human and yeast proteasomes, likely due to the insufficient resolution in these regions. Here, we show that, collectively, the conserved linker proline residues in Rpt2, Rpt3, and Rpt5 are important for proper proteasome base assembly in *S. cerevisiae*. Furthermore, we provide evidence for the role of Hsp42 in promoting base assembly in yeast expressing proline-to-alanine mutations in both Rpt2 and Rpt5 by suppression of the aggregation of these subunits.

## Results

### Importance of conserved Rpt linker prolines for proteasome assembly

Based on phylogenetic analysis of proteasomal ATPases from deeply divergent eukaryotes, we found that the N-domain linker proline is strictly conserved in Rpt3 and Rpt5, highly conserved in Rpt2, and only loosely conserved in Rpt1 (Figure 2A; Table S1). Since Rpt2, Rpt3, and Rpt5 belong to distinct heterodimer pairs, this finding is consistent with the hypothesis that the conserved proline residue in these subunits allows the *cis* peptide conformation. This should kink the CC-OB linker and facilitate helix interaction and CC formation with the *trans* Rpt partner (14,23).

**Figure 2.**
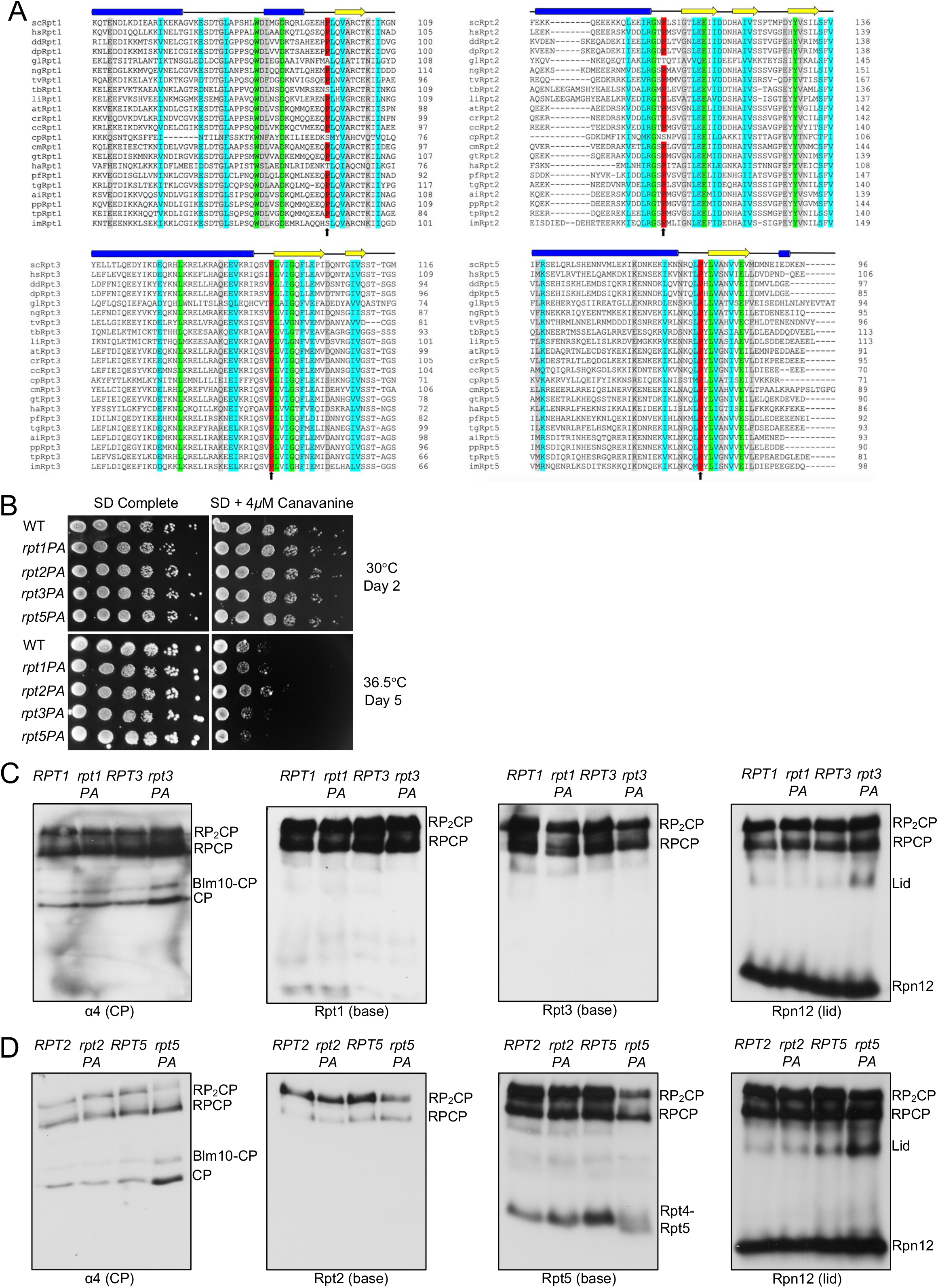
Rpt P-to-A mutants have no growth defects but minor proteasome assembly defects. (A) Sequence alignment of Rpt1, Rpt2, Rpt3, and Rpt5 depicting conservation of the proline residue in diverse eukaryotic species. Structural domains above the alignments are based on secondary structure predictions (via PSIPRED) of *S. cerevisiae* subunits. Sequence alignments conducted with EMBOSS (EMBL-EBI) alignment tool. (Green: complete conservation of amino acid; cyan: conservation between groups with strongly similar properties; gray: conservation between groups with weakly similar properties). Species selected from all six recognized eukaryotic supergroups See Table S1 for species abbreviations (B) Growth assays of single P-to-A mutants. No obvious growth defects are evident. Yeast cultures were subjected to six-fold serial dilutions and spotted on the indicated plates. (C) Visualization of proteasomes by immunoblot analyses of yeast *RPT1/rpt1-P96A* and *RPT3/rpt3-P93A* whole cell extracts separated by nondenaturing PAGE. Strains were grown in selective defined media at 30°C to log phase. RP_2_-CP and RP-CP are doubly and singly capped 26S proteasomes. (D) Visualization of proteasomes by immunoblot analyses of yeast *RPT2/rpt2-P103A* and *RPT5/rpt5-P76A* whole cell extracts separated by nondenaturing PAGE. Strains were grown as in (C).

To determine the importance of these prolines *in cellulo*, we made Pro-to-Ala substitutions at Rpt1-P96, Rpt2-P103, Rpt3-P93, and Rpt5-P76 and determined their impact on yeast growth. Perhaps surprisingly, none of the resulting single mutants showed obvious growth defects compared to wild-type (WT) cells even under proteotoxic stress conditions (Figure 2B). To investigate if these mutations affect proteasome assembly, we subjected whole cell lysates from these strains to native gel immunoblot analyses. Despite their lack of obvious growth impairment, the *rpt3-P93A* and *rpt5-P76A* mutants exhibited detectable RP base assembly defects. Assembly was most strongly retarded in the *rpt5-P76A* mutant based on a decrease in doubly-capped 26S proteasomes, excess accumulation of free CP and Blm10-CP (Blm10 is an alternative CP regulator), and increased levels of free lid subcomplex (Figure 2C and 2D).

Consistent with the effects of the above Pro-to-Ala mutations on proteasome base assembly, they also caused synthetic growth defects at elevated temperature when combined with *hsm3*Δ; loss of Hsm3 has the strongest effect on growth of any single base assembly chaperone mutant (Table 1; Figure S1) (5–9). The *rpt2-P103A* and *rpt5-P76A* alleles also displayed synthetic defects with *nas2* Δ and *nas6* Δ , respectively. The latter each lack a base assembly chaperone that promotes assembly of an Rpt heterodimer not affected directly by the respective Rpt Pro-to-Ala mutation; thus, two different base assembly modules are impacted in these mutant combinations, possibly accounting for the synthetic effects on growth. By contrast, no additional growth defects were observed when these Rpt mutations were crossed into yeast strains with the CP assembly chaperone deletions *pba1*Δ or *pba4*Δ. Notably, *rpt1-P96A*, which affects the ATPase subunit with the least conserved linker proline, did not exhibit synthetic defects with any tested base assembly chaperone deletions (Table 1; Figure S1).

**Table 1.**
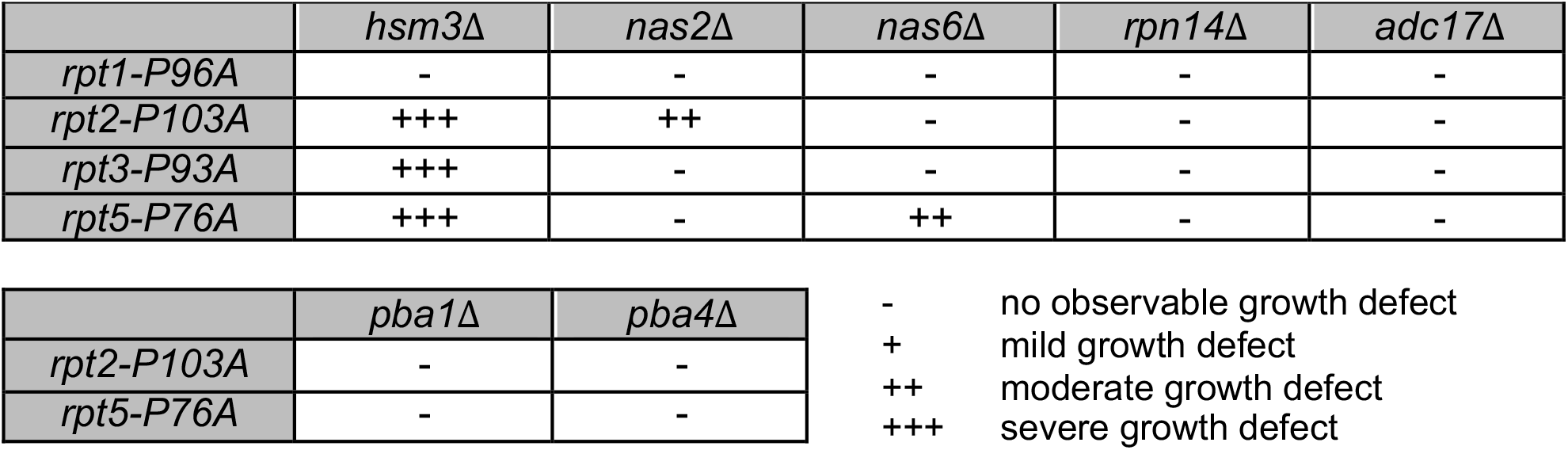
Synthetic genetic interactions between *rpt1-P96A*, *rpt2-P103A*, *rpt3-P93A*, or *rpt5-P76A* with base and CP assembly chaperone gene deletions Yeast growth was analyzed by streak tests on YPD at 36°C. Growth defect noted is relative to congenic yeast expressing the WT *RPT* alleles in strains with the indicated assembly chaperone gene deletions.

|  | *hsm3Δ* | *nas2Δ* | *nas6Δ* | *rpn14Δ* | *adc17Δ* |
| --- | --- | --- | --- | --- | --- |
| *rpt1-P96A* | - | - | - | - | - |
| *rpt2-P103A* | +++ | ++ | - | - | - |
| *rpt3-P93A* | +++ | - | - | - | - |
| *rpt5-P76A* | +++ | - | ++ | - | - |

|  | *pba1Δ* | *pba4Δ* |
| --- | --- | --- |
| *rpt2-P103A* | - | - |
| *rpt5-P76A* | - | - |

- no observable growth defect + mild growth defect ++ moderate growth defect +++ severe growth defect

### The Rpt5 linker proline is important for Rpt5 stability and solubility

Of the four individual Pro-to-Ala mutants analyzed, *rpt5-P76A* showed the strongest proteasome assembly defects, but the mutant still appeared to grow normally. We investigated whether *rpt5-P76A* exhibited a synthetic defect when combined with a deletion of the proteasome transcription factor gene *RPN4*. Rpn4 is required for normal levels of proteasome subunit transcription and is upregulated when proteasome activity is reduced (25,26). Indeed, deletion of *RPN4* resulted in substantial synthetic growth defects with *rpt5-P76A* (Figure 3A). This was not observed when *rpn4*Δ was combined with *rpt1-P96A*, *rpt2-P103A*, or *rpt3-P93A* (Figure S2A). The growth defects of *rpn4*Δ *rpt5-P76A* were paralleled by strongly reduced 26S proteasome formation *in vivo* (Figure S2B). Quantitative RT-PCR analysis also revealed slight but consistent increases in proteasome subunit transcript levels in *rpt5-P76A* relative to *RPT5* cells, as expected if Rpn4-induced transcription was partially compensating for reduced RP base assembly in the mutant (Figure 3B). When we made Ala substitutions in the two residues flanking Rpt5-P76 (Rpt5-L75A and Rpt5-Y77A), little or no synthetic defect was seen with *RPN4* deletion, demonstrating the specificity of the *rpn4*Δ *rpt5-P76A* interaction (Figure 3C).

**Figure 3.**
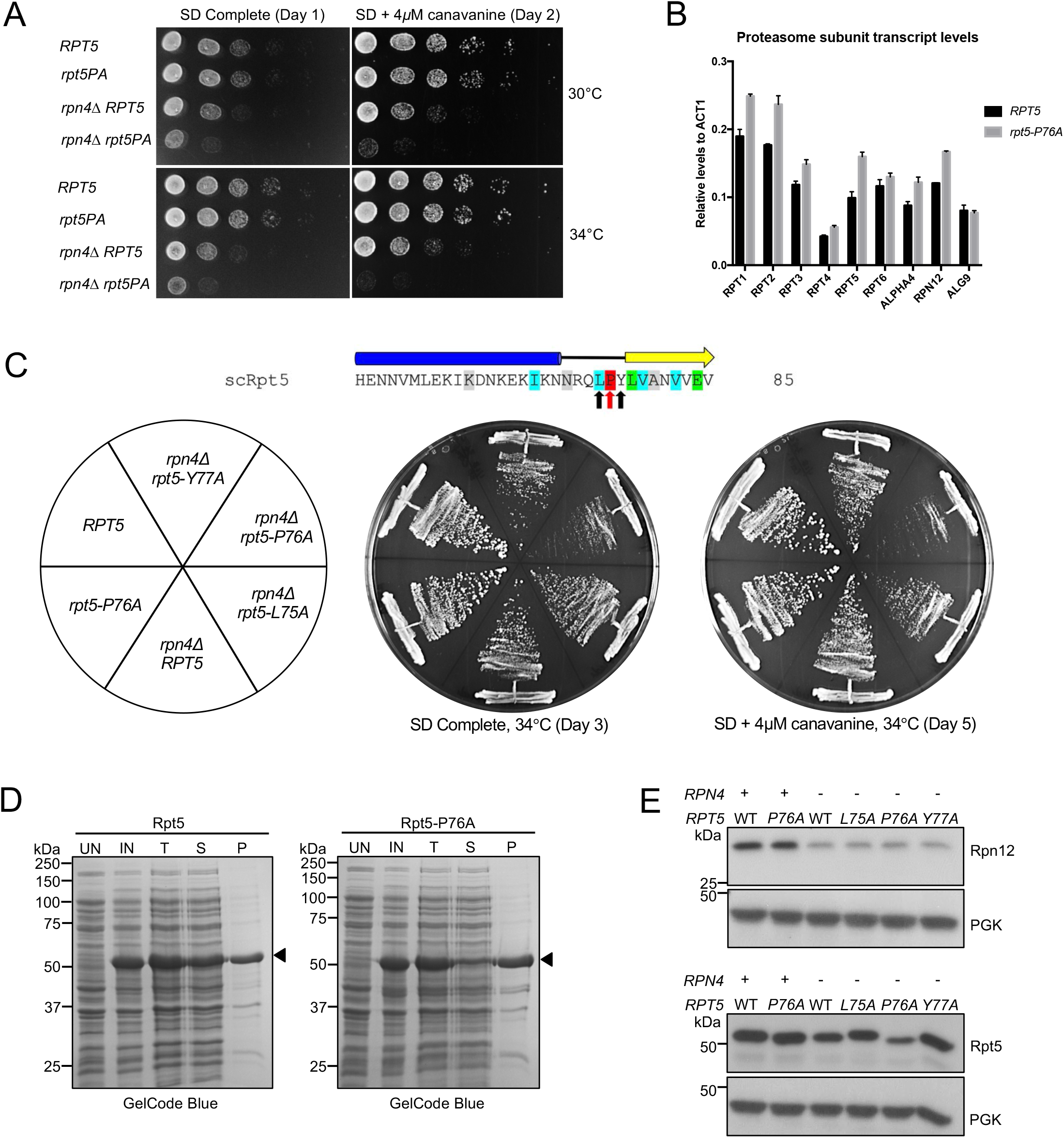
Rpt5-P76A has unique defects among the P-to-A linker mutants. (A) Synthetic growth defects of an *rpt5-P76A rpn4*Δ double mutant. Cells were spotted as in Fig. 2B. (B) Transcript levels for all proteasome subunits are consistently higher in *rpt5-P76A* relative to *RPT5* cells. *ALG9* serves as an internal control. (mean ± SD; n=3, technical replicates). (C) Growth assays of strains streaked on the indicated plates. A synthetic defect with *rpn4*Δ is seen with *rpt5-P76A* but is not observed with the flanking *rpt5-L75A* and *rpt5-Y77A* mutations. In the schematic above, Rpt5-P76 is indicated with a red arrow, while the flanking mutated residues are indicated with black arrows. (D) A higher fraction of bacterially expressed recombinant Rpt5-P76A is insoluble compared to WT Rpt5. Arrowheads denote WT Rpt5 or Rpt5-P76A protein. UN, Uninduced; IN, Induced; T, Total protein; S, Supernatant; P, Pellet. (E) Steady-state levels of soluble Rpt5-P76A are lower relative to WT Rpt5 in an *rpn4*Δ background. Yeast strains were grown in YPD at 30°C to log phase. Phosphoglycerate kinase (PGK) served as a loading control.

Next, we expressed recombinant WT Rpt5 and mutant Rpt5-P76A in *Escherichia coli* and determined the solubility of these proteins via a pelleting assay. For this, we lysed bacterial cells expressing each protein under nondenaturing conditions and subjected the lysates to centrifugation. The supernatant (S) fraction contained soluble proteins and the pellet (P) fraction included aggregated proteins. Rpt5-P76A had a much higher propensity to aggregate relative to WT Rpt5, as most of the mutant protein was in the pellet (Figure 3D). This observation was paralleled by the finding that steady-state levels of soluble Rpt5 were lower in *rpn4*Δ *rpt5-P76A* relative to *rpn4*Δ *RPT5* yeast (Figure 3E). These data suggest that Rpt5-P76A is prone to misfolding and aggregation. Rpt5-P76A aggregation is associated with proteasome assembly defects in mutant yeast cells and consequently, upregulation of proteasome subunit genes via Rpn4 to compensate for the depletion of the compromised mutant subunit.

### Double *rpt2-P103A rpt5-P76A* mutant has synthetic assembly and growth defects

In the archaeal PAN ATPase, the single PAN-P91A mutation disrupts all subunits of the hexamer but in particular the three “*cis*” subunits, resulting in a severe defect in ring assembly (14). We investigated the effect on yeast growth of proline-to-alanine mutations in pairs of Rpt subunits. Out of the six possible double mutant combinations, we found that only one, *rpt2-P103A rpt5-P76A (rpt2,5PA)*, resulted in a growth defect, which was severely exacerbated at elevated temperature (Figure 4A; Figure S3). To determine the specificity of the negative synthetic interaction between *rpt2-P103A* and *rpt5-P76A*, we tested if double mutant combinations with mutations on both flanking residues of Rpt2-P103 and Rpt5-P76 result in similar growth defect as *rpt2,5PA*. Out of all double mutant combinations tested, only *rpt2-L104A rpt5-P76A* displayed a moderate growth defect at elevated temperature on SD+4*μ*M canavanine plate, although the defect was not nearly as severe as that of *rpt2,5PA* (Figure 4B).

**Figure 4.**
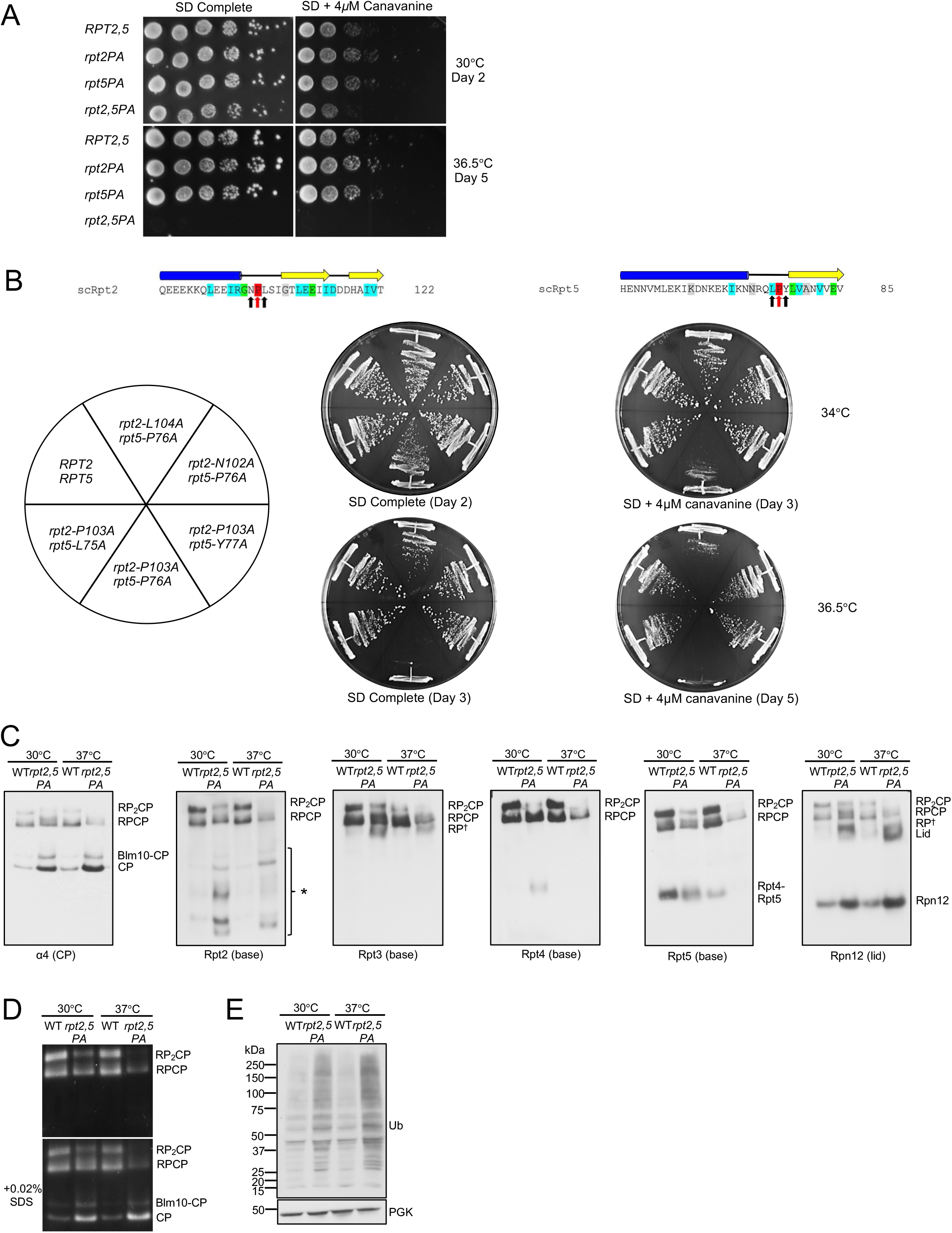
The *rpt2-P103A rpt5-P76A* double mutant has a strong synthetic growth defect. (A) Growth assays of the *rpt2-P103A rpt5-P76A* (*rpt2,5PA)* double mutant compared to WT and single mutant cells. Cultures were spotted as in Figure 2B. (B) Growth assays highlighting the specificity of the strong *rpt2,5PA* growth defect. A mild synthetic growth defect was also observed in *rpt2-L104A rpt5-P76A* cells. (C) Visualization of proteasome complexes by immunoblot analyses of yeast *rpt2,5PA* whole cell extracts separated by nondenaturing PAGE. Defects are more severe in cells grown at elevated temperature. Yeast strains were grown in YPD to log phase at the indicated temperatures. *, Rpt2-containing subcomplexes; RP^†^, RP or RP-like complex. (D) Suc-LLVY-AMC substrate overlay assay depicts lower overall proteasome activity in the *rpt2,5PA* mutant. SDS addition to the gel allows visualization of free CP activity. (E) Western blot analysis of yeast whole cell lysate resolved on a denaturing gel shows accumulation of ubiquitin-protein conjugates in *rpt2,5PA* mutant. Anti-PGK blotting used to show similar sample loading.

The *rpt2,5PA* mutant had a pronounced proteasome assembly defect characterized by accumulation of free CP, Blm10-CP, and lid subcomplex and decreased levels of singly (RPCP) and especially doubly capped (RP_2_CP) 26S proteasomes (Figure 4C). In addition to these species, the double mutant also accumulated a number of novel Rpt2-containing subcomplexes, which might be dead-end complexes (Figure 4C). Consistent with the decrease in level of full proteasomes, the mutant cells exhibited lower total proteasomal peptidase activity (Figure 4D) and an increased accumulation of cellular ubiquitin conjugates (Figure 4E). Defects in proteasome assembly and activity in the double mutant were worse at elevated temperature (Figure 4C-E).

### Rpt5-PA ubiquitination and mutant E3 Not4-L35A suppression of *rpt2,5PA*

Analyses of steady-state levels of proteasome subunits revealed that the overall levels of subunits in *rpt2,5PA* did not decrease at either permissive or non-permissive temperature (Figure 5A). In fact, overall levels of subunits increased despite the strong reduction in fully formed proteasomes in the mutant strain, hinting at a possible defect in elimination of defective proteasome subunits/sub-complexes. Indeed, we observed an accumulation of high molecular weight (HMW) Rpt5-containing species that could be ubiquitinated forms of Rpt5-PA (Figure 5A). To confirm this, we conducted ubiquitin pulldown assays and found higher levels of ubiquitinated Rpt5 in the *rpt2,5PA* mutant compared to the WT strain (Figure 5B). It has been previously reported that Rpt ring assembly is regulated through selective ubiquitination of Rpt5 by the E3 ligase Not4 (27). When ubiquitination sites on Rpt5 are exposed during Rpt ring assembly due to the absence of Hsm3 and Nas2 binding or during defective base assembly, Rpt5 is selectively ubiquitinated and ring assembly is inhibited (27). We speculated that Not4 similarly inhibits ATPase ring assembly in *rpt2,5PA* cells due to the presence of these mutant Rpt5 species. Indeed, when *NOT4* was replaced with the catalytic mutant *not4-L35A*, the growth defect of *rpt2,5PA* was partially suppressed (Figure 5C). We have also found that Not4-regulation of proteasome assembly is likely specific to RP base mutants as *not4-L35A* did not suppress the temperature-sensitivity of either *pre9*Δ (CP subunit) or *sem1*Δ (lid subunit) (Figure S4).

**Figure 5.**
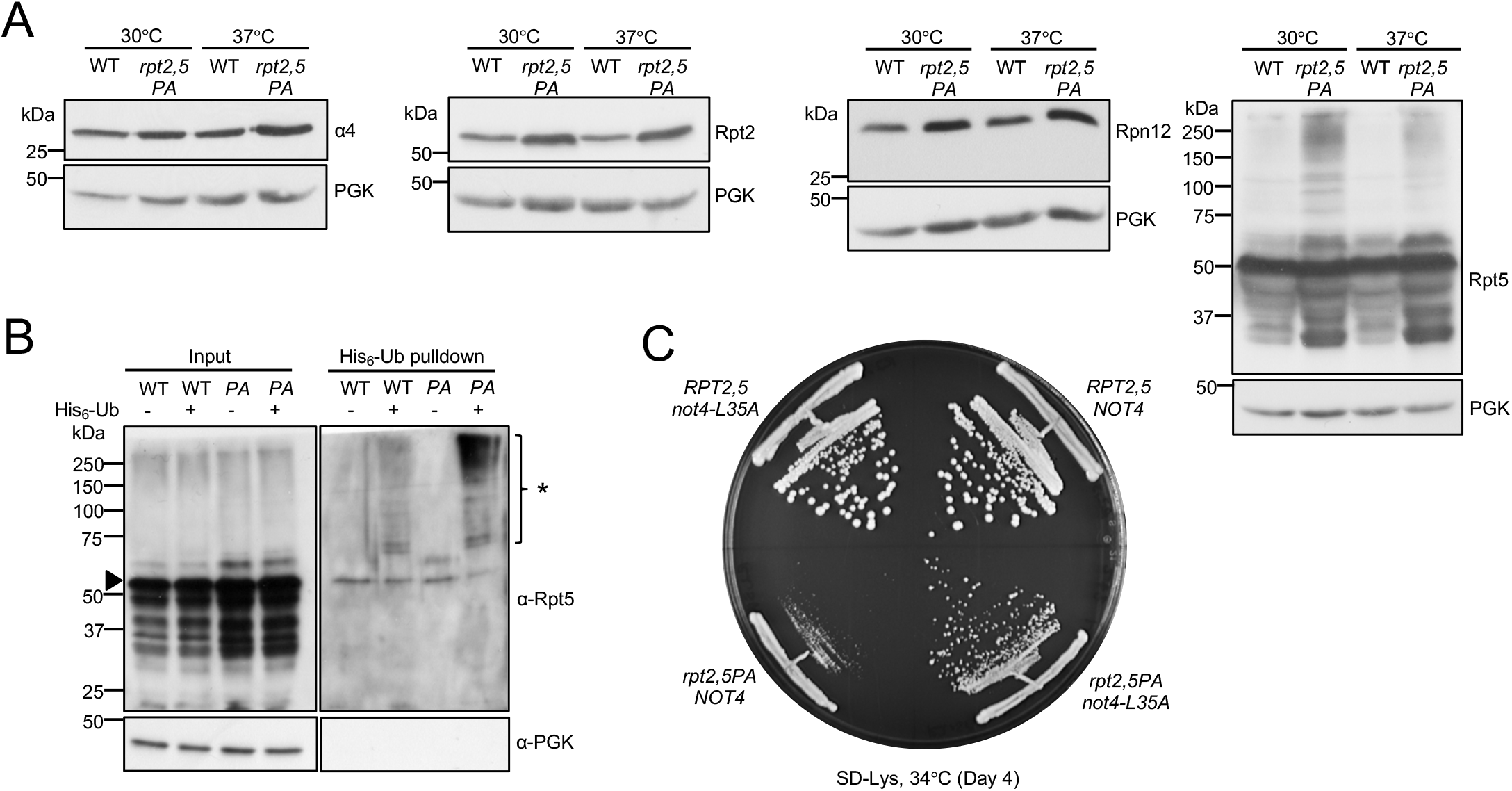
Increased ubiquitination of Rpt5 in *rpt2,5PA* and partial suppression of *rpt2,5PA* growth defect by *not4-L35A* E3 mutation. (A) Increased proteasome subunit steady-state levels (left three panels) and accumulation of high molecular-mass species of Rpt5-P76A (rightmost panel) in *rpt2,5PA* cells. Anti-PGK blotting used as a control for sample loading. Yeast strains were grown in YPD at the indicated temperatures to log phase. (B) Purification of His6-tagged ubiquitin conjugates using a Ni-NTA resin reveals higher levels of ubiquitinated Rpt5 in *rpt2,5PA* (PA) cells, especially at non-permissive temperature. Eluted proteins were resolved by SDS-PAGE and immunoblotted with anti-Rpt5 antibodies. Arrowhead denotes unmodified Rpt5 band. *, ubiquitinated Rpt5 species. (C) Expression of Not4-L35A partially suppresses the *rpt2,5PA* growth defect. Wild-type Not4 or Not4-L35A was expressed from a low-copy pRS317 plasmid under its native promoter in WT *RPT2,5* or mutant *rpt2,5PA* strains with the chromosomal *NOT4* gene deleted.

### Hsp42-mediated PQC regulates cell fitness and aggregation of Rpt2 and Rpt5

Because overall levels of Rpt subunits in *rpt2,5PA* did not decrease even at elevated temperature (Figure 5A), and levels of full proteasomes and soluble Rpt intermediates/subcomplexes were further reduced at elevated temperature (Figure 4C), we speculated that Rpt subunits in this mutant have the propensity to be sequestered either as storage for future use or as insoluble aggregates for degradation or elimination via mother cell retention. We conducted aggregation assays using yeast whole cell lysates to determine if the subunits form insoluble aggregates at elevated temperature (Figure 6A). We found that the double mutant had a higher pellet (2xP) to supernatant (S) ratio for Rpt2 and Rpt5 subunits relative to wild-type, suggesting that these subunits aggregate in the mutant (Figure 6B). Unlike Rpt2 and Rpt5, aggregation was not as prominent in CP subunit (α4) and was absent in lid subunit (Rpn12) in the mutant strain (Figure S5A). We have also found that the Rpt3 subunit aggregated in the mutant but Rpt4 did not (Figure S5A). However, overall levels of Rpt4 seemed to be lower in the mutant in samples collected at saturation phase, suggesting that Rpt4 expression might be suppressed or that it is selectively degraded (Figure S5A). We believe that the smears observed predominantly above Rpt2 and Rpt3 monomers in (T) and (2xP) in Rpt2 and Rpt3 immunoblots are primarily SDS-resistant aggregates that are recognized non-specifically by Rpt2 and Rpt3 antibodies. Ubiquitin pulldown assay further suggested that the smear above Rpt3 monomer also contained a small fraction of ubiquitinated Rpt3 although the levels were similar in both wild-type and mutant strains (Figure S5B).

**Figure 6.**
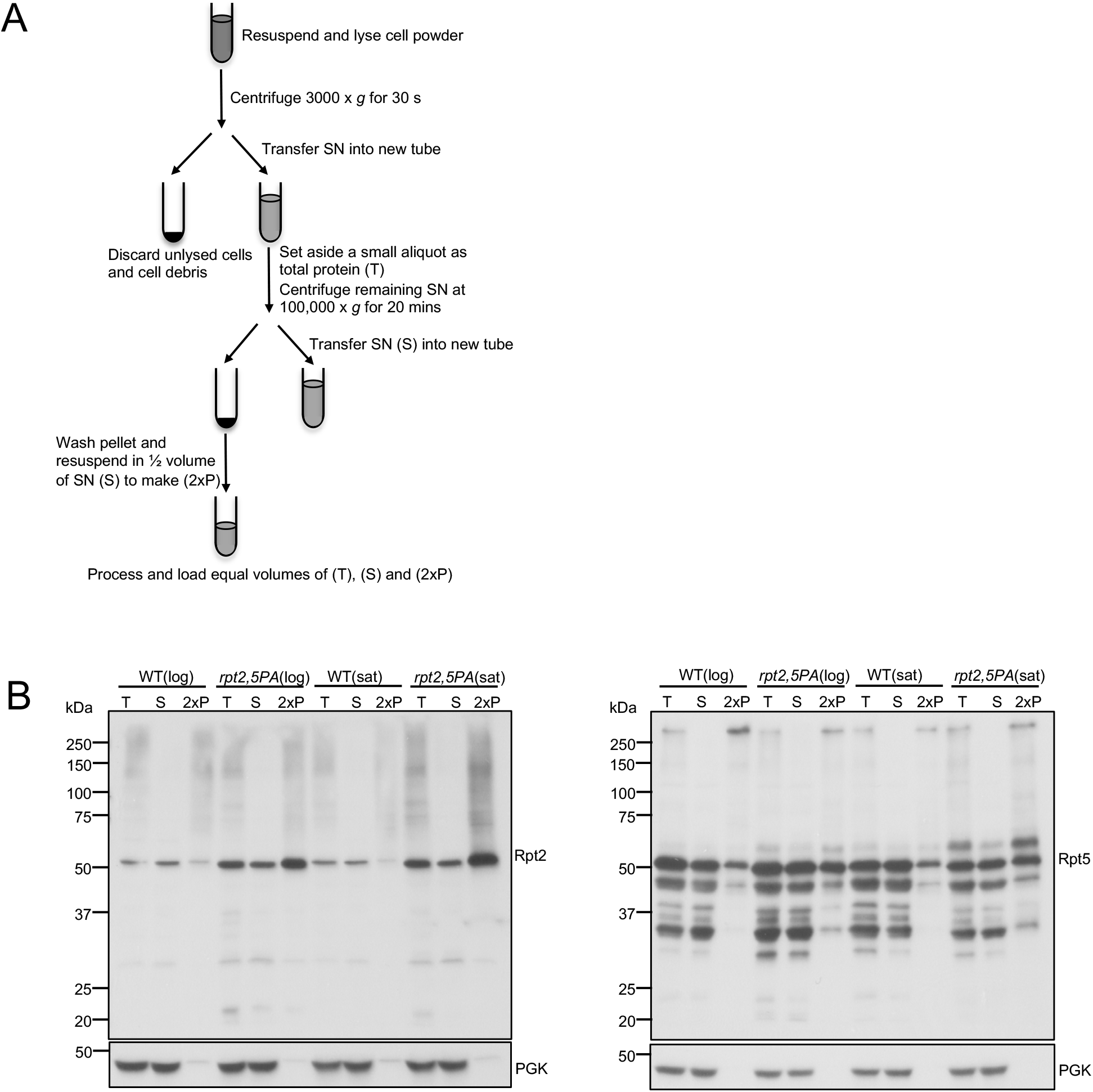
Rpt2 and Rpt5 subunits in *rpt2,5PA* cells are prone to aggregation. (A) Aggregation assay workflow. Yeast strains were grown in synthetic defined medium (with casamino acids) at 37°C. (B) Increased Rpt2 and Rpt5 aggregation at high temperature as seen by an increase of these proteins in pellet (P) fraction compared to supernatant (S) in extracts from *rpt2,5PA* cells. Total protein, T. Anti-PGK blotting used as a control for relative (soluble) protein loading.

A previous study showed that a lid mutant (*rpn5*Δ*C*) forms aggregates at elevated temperature and is regulated by the PQC machinery via a group of heat-shock proteins, primarily Hsp42 (28). Deletion of *HSP42* in *rpn5*Δ*C* prevents sequestration of Rpn5ΔC and allows more Rpn5ΔC to assemble into full proteasomes, thereby strongly suppressing growth defect of the *rpn5* Δ *C* mutant (28). We wanted to determine if *rpt2,5PA* is similarly regulated by this PQC machinery. We found that *hsp42*Δ partially suppressed the *rpt2,5PA* growth defect, albeit to a much lower extent compared with *rpn5*Δ*C* (Figure 7A) (28). Although proteasome activity assay revealed that the *hsp42* deletion in the double mutant showed little to no suppression in proteasome assembly (Figure 7B), an aggregation assay showed a suppression in the aggregation of Rpt2 and Rpt5 subunits, consistent with partial growth rescue observed (Figure 7C). This finding suggests that the Hsp42-mediated PQC is responsible, at least in part, for the regulation of proteasome base subunits. Interestingly, we found that *hsp42* deletion also partially rescued temperature-sensitivity of other base (*cim3-1* and *rpt4-G106D*) and CP (*pre9*Δ) mutants (Figure S6).

**Figure 7.**
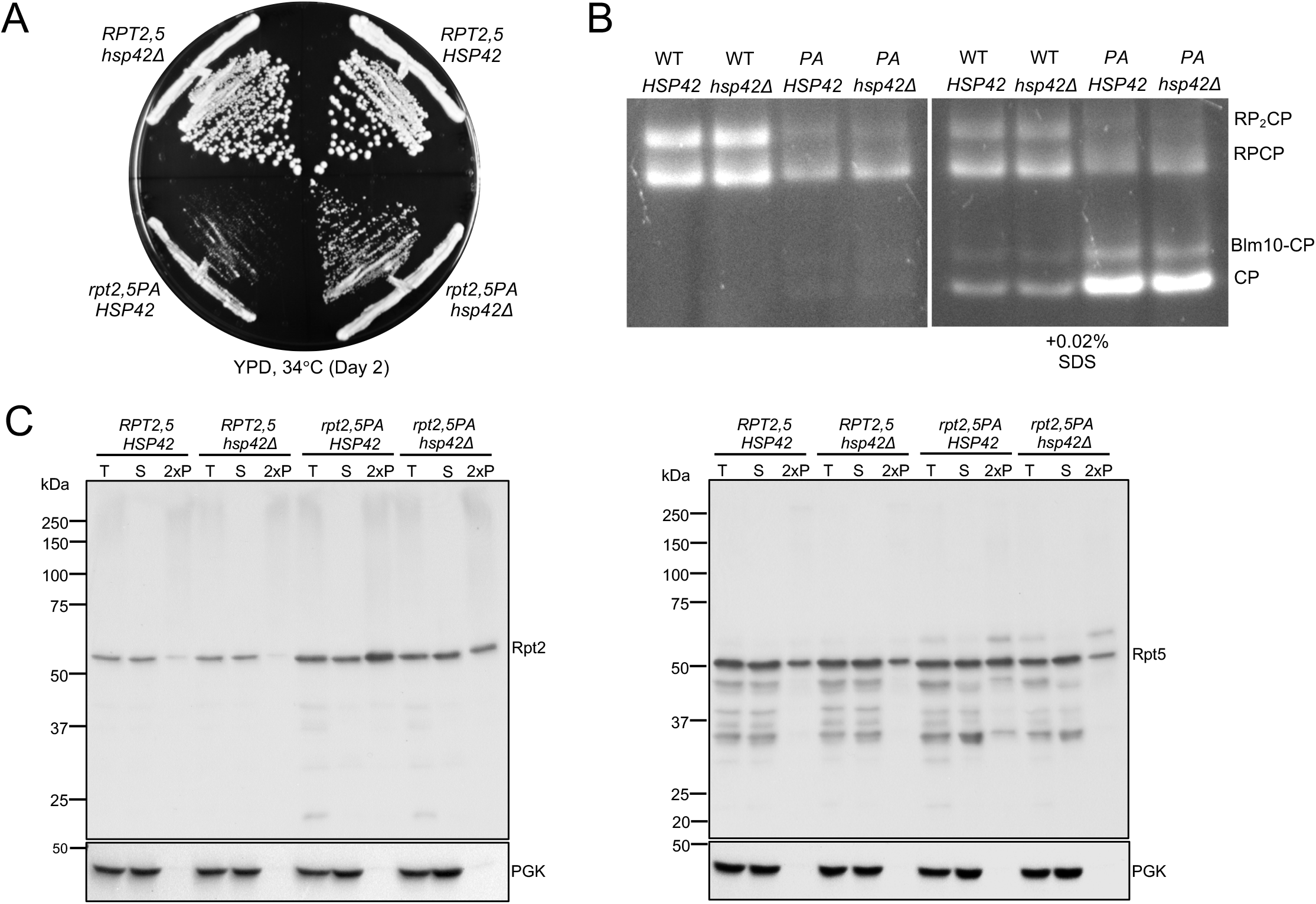
Hsp42 promotes proteasome subunit aggregation in *rpt2,5PA* cells. (A) *hsp42*Δ partially suppresses the growth defect of *rpt2,5PA* cells. (B) Suc-LLVY-AMC proteasome activity assay indicates *hsp42* Δ may very weakly suppress the proteasome assembly defect of *rpt2,5PA*. Yeast strains were grown in YPD at 37°C to log phase. (C) Aggregation assay reveals that *HSP42* deletion in *rpt2,5PA* partially suppresses aggregation of Rpt2 and Rpt5. Yeast strains were grown in YPD at 37°C to log phase.

## Discussion

Here we have shown that the conserved Rpt linker prolines promote eukaryotic 26S proteasome base assembly, most likely by facilitating specific pairwise ATPase heterodimerization. Based on structural data from their prokaryotic counterparts, this is potentially due to the enhanced ability of proline residues to form *cis* peptide bonds; this would create a kink in the linker that allows the upstream helical domain of the subunit to form a coiled coil more readily with its (*trans*) ATPase partner (14,23). Interestingly, recent work has suggested another mechanism for promoting specific Rpt heterodimer interaction that involves pausing of the ribosome during translation of the Rpt1 and Rpt2 nascent complexes, to allow their co-translational assembly (29). The disordered N-terminal segment of Rpt1 was shown to be important for pausing, and the Not1 subunit of the Ccr4-Not complex (which includes Not4) participates in colocalizing the stalled translation complexes. All three Rpt heterodimers also interact with dedicated (and non-homologous) RP assembly chaperones (RACs); for the Rpt3-Rpt6 dimer, three different RACs help promote its assembly (5–10). Hence, eukaryotes have evolved multiple mechanisms to increase the assembly efficiency and fidelity of the proteasomal heterohexameric ATPase ring.

### Single linker Pro mutations are tolerated to varying degrees

Yeast cells show a surprisingly high tolerance to single Rpt proline-to-alanine mutations based on growth assay analysis (Figure 2B). For most of the mutants, assembly appears normal or nearly so; despite the lack of a growth defect, *rpt5-P76A* displays a modest proteasome assembly defect (Figure 2D). The lower solubility of Rpt5-P76A is reminiscent of the aggregation seen with the homologous archaeal PAN-P91A mutation (14). Assembly of an Rpt heterohexamer in eukaryotes, rather than a homohexamer, may limit growth and assembly defects due to single P-to-A mutations; the introduction of a single PAN-P91A mutation effectively disrupts all three “*cis*” subunits of the homohexamer. Consistent with this, base assembly is far more severely disrupted in the double *rpt2-P103A rpt5-P76A* mutant (Figure 4).

The presence of dedicated base assembly chaperones may also suppress the Rpt P-to-A mutations. In support of this possibility, the *rpt2-P103A*, *rpt3-P93A*, and *rpt5-P76A* mutations have strong synthetic defects with *hsm3*Δ and with other select RAC gene deletions (Table 1; Figure S1). By contrast, *rpt1-P96A* did not display synthetic defects with any RAC deletions. Rpt2, Rpt3, and Rpt5 belong to distinct dimer pairs in early base assembly, and their linker prolines are more highly conserved than in Rpt1. Our findings are consistent with the hypothesis that Rpt2, Rpt3, and Rpt5 are the “*cis*” subunits in the eukaryotic Rpt ring, a conclusion that still awaits structural confirmation.

### Synthetic defects in *rpt2,5PA* mutant linked to severely impaired base assembly

The *rpt2,5PA* strain is the only double P-to-A mutant that displayed strong growth and proteasome assembly defects (Figure 4). We speculate that the peptide bonds preceding these prolines have to be in a *cis* conformation for Rpt2 and Rpt5 to associate efficiently with their partner ATPase subunits during assembly into higher-order base subcomplexes. Rpt2-P103A and Rpt5-P76A mutations may, for example, inhibit coiled-coil formation between the correct pairs of Rpt subunits and thereby disrupt proper base assembly. The tendency of Rpt5-P76A to aggregate may further enhance base assembly defects in the double mutant. A decreased ability of Rpt2- and Rpt5-containing complexes to properly associate could therefore lead to the formation of Rpt2-containing dead-end complexes or assembly intermediates, which could explain the presence of several Rpt2-containing complexes that are not found in WT strains based on our native immunoblot analyses (Figure 4C). The reduced accumulation of these unique complexes without an increase in higher-order complexes and full proteasomes at elevated temperature further suggests that these complexes are unstable and subsequently degraded and/or sequestered into aggregates.

High levels of ubiquitinated Rpt5-P76A also accumulate in the *rpt2,5PA* strain, suggesting that the misassembled ATPase subunit is marked for ubiquitin-dependent degradation. Expression of the Not4-L35A ubiquitin ligase catalytic mutant partially suppresses the growth defect of the *rpt2,5PA* mutant, consistent with the presence of a base assembly defect and a role for Not4 in regulating base assembly in this mutant (Figure 5) (27). The modest suppression observed relative to the base assembly mutants studied by Fu et al. could be due to base assembly defects in *rpt2,5PA* cells that go beyond simply exposing Not4 ubiquitination sites on Rpt5; inhibition of Not4 catalytic ligase activity might therefore be insufficient to substantially promote base assembly in this mutant.

### Hsp42 participates in the aggregation of proteasome base subunits in *rpt2,5PA* cells

We found that Rpt2 and Rpt5 subunits aggregate in the *rpt2,5PA* double mutant (Figure 6B). We attempted to tag the N-termini of Rpt2 and Rpt5 with GFP to track these aggregates via microscopy, but the resulting strains were inviable. Nevertheless, we found that deletion of *HSP42* partially suppressed growth defects of *rpt2,5PA*, although not to the extent seen with *rpn5*Δ*C* (Figure 7A) (28). The (partial) suppression of both the growth defect and Rpt2/Rpt5 subunit aggregation by *hsp42*Δ suggests that the Hsp42-based PQC machinery is important for regulating proteasome sequestration or PQC in distinct mutants of the proteasome, not just RP lid mutants (Figure 7A and 7C). Indeed, our data indicate that Hsp42 is a general regulator of proteasome assembly, as revealed by the ability of *hsp42*Δ to also suppress other RP base (*cim3-1* and *rpt4-G106D*) mutants and a CP (*pre9*Δ) mutant (Figure S6). Interestingly, *hsp42*Δ did not suppress another lid mutant, *sem1*Δ, possibly because Sem1 is involved not only in proteasome assembly (30) but also in the functioning of the mature 26S proteasome (31–33) as well as that of other protein complexes (34,35).

In summary, our data are consistent with the hypothesis that the highly conserved Rpt linker prolines promote formation of *cis* peptide bonds specifically in one subunit of each eukaryotic Rpt heterodimer, which facilitates their dimerization with the correct “*trans*” subunits, presumably through enhanced coiled-coil formation. Yeast cells have multiple mechanisms that allow them to tolerate mutations in the highly conserved linker prolines of the three predicted “*cis*” subunits Rpt2, Rpt3, and Rpt5. These include the Rpn4-dependent transcriptional feedback loop and Rpt heterodimer-specific assembly chaperones. On the other hand, PQC mechanisms that result in ubiquitination and degradation of mutant or misassembled subunits of the proteasome or the sequestration of aberrant assembly intermediates into Hsp42-dependent aggregates such as IPODs enhance growth deficiencies of these mutants. Together with the recent description of co-translational assembly of Rpt heterodimers, our results point to the importance of multiple mechanisms, which are likely intertwined, to ensure efficient and high-fidelity assembly of the eukaryotic 26S proteasome.

## Experimental procedures

### Yeast strains

Yeast strains were made following standard procedures (36). Yeast haploid strains with WT or proline-to-alanine Rpt subunits expressed from low-copy plasmids and their native promoters were created in strains with the corresponding chromosomal copy or copies replaced with a *HIS3* cassette as described previously (4). Because all *RPT* genes are essential, the parental strains all initially had the relevant WT *RPT* gene(s) on plasmids bearing a *URA3* selectable marker. Strains were then transformed with plasmids carrying either a *TRP1* or *LEU2* selectable marker and expressing either WT or mutant *rpt* alleles. The resulting strains were then cured of the original *URA3* plasmid by counterselection on 5-fluoroorotic acid (5-FOA). The list of yeast strains and plasmids used can be found in Tables S2 and S3, respectively.

### Yeast growth assays

Yeast strains were grown in YPD rich medium or selective defined media to saturation overnight. The next day, strains were diluted in sterile water to 0.2 OD_600_ units in a final volume of 1 mL. Samples were then spotted in a six-fold dilution series on the appropriate plates and incubated at various temperatures, and growth was monitored over several days.

### Nondenaturing gel analyses of proteasomes in whole cell extracts

Yeast extracts for nondenaturing gel analyses were prepared as previously described with slight modifications (37). Yeast cultures were grown in YPD or selective defined media overnight. The next day, cultures were diluted to OD_600_=0.2 in YPD or selective defined media and grown to mid-log phase (unless otherwise stated), washed with ice-cold sterile water, and subsequently frozen in liquid nitrogen and stored at −80°C. Frozen cells in liquid N_2_ were ground using a mortar and pestle until a fine powder formed. The resulting powder was collected in a pre-chilled tube and incubated in proteasome extraction buffer (50 mM Tris-HCl pH 7.5, 5 mM MgCl_2_, 10% glycerol, 5 mM ATP) for 10 min with occasional vortexing. Samples were then centrifuged at 22,000 x *g* for 10 min to remove unlysed cells and cell debris. The resulting supernatants were collected and protein concentration was determined using the BCA assay conducted according to manufacturer’s instructions (Thermo Fisher Scientific). 50 *μ*g samples were run onto 4% nondenaturing gels. Gels were either overlayed with a fluorogenic substrate, Suc-LLVY-AMC (Sigma-Aldrich) or were used in immunoblot analyses. Details of the experimental procedures for the in-gel peptidase assay are as described (38). For analyses of steady-state levels of soluble protein from these lysates (as in Figure 3E), 10 *μ*g of each supernatant were run in denaturing SDS gels and subjected to immunoblotting.

### Denaturing gel analyses of overall levels of proteins in yeast extracts

Yeast extracts for denaturing gel analyses were prepared as previously described with slight modifications (39). Yeast cultures were grown as above (unless otherwise noted). 2.5 OD_600_ units of cells were harvested by centrifugation and washed with ice-cold sterile water. Samples were then resuspended in 200 *μ*L sterile water followed by the addition of 200 *μ*L 0.2 M NaOH and incubated at room temperature for 5 min with occasional vortexing. Cells were pelleted at 10,000 x *g* for 1 min, and supernatants were discarded. Pelleted cells were resuspended in 1X SDS-PAGE sample buffer containing 4% b-mercaptoethanol (BME) and heated at 100°C for 5 min followed by centrifugation at 10,000 x *g* for 1 min; 10-15 *μ*L of the supernatants were resolved in discontinuous SDS gels and subjected to immunoblot analyses.

### Aggregation assay of recombinant 6His-Rpt5 expressed in E. coli

Competent Rosetta DE3 cells transformed with either pET15b-6His-Rpt5 or pET15b-6His-Rpt5-P76A plasmid were grown in LB + 100 *μ*g/mL ampicillin (Amp) media overnight at 37°C. Cultures were diluted 1:100 in fresh LB + Amp medium and grown to OD_600_=0.6-0.8. One OD_600_ unit of culture was removed as uninduced (UN) sample. Cultures were then induced with 0.2 mM isopropyl β-D-1-thiogalactopyranoside (IPTG) (final) and grown at 16°C overnight. One OD_600_ unit of culture was harvested as induced (IN) sample. Another 1.5 mL aliquot from each culture was harvested for aggregation assays and resuspended in 700 *μ*L lysis buffer (50 mM Tris-HCl pH 7.5, 150 mM NaCl, 10% glycerol, 100 *μ*g/mL lysozyme, 1 mM phenylmethylsulfonyl fluoride (PMSF)) and incubated at 4°C for 30 min. Samples were then sonicated 6 ×10 s with 10 s incubations on ice between each sonication round; 100 *μ*L aliquot from each sample was transferred into a new tube and represented total protein (T). The remaining samples were centrifuged at 21,000 x *g* for 5 min at 4°C. The supernatant (S) was transferred into a new tube. The pellet (P) was washed once with 600 *μ* L lysis buffer, re-centrifuged as above; lysis buffer was removed and the pellet was resuspended in 600 *μ*L lysis buffer. UN and IN cell pellets were resuspended in 150 *μ*L of 1X SDS sample buffer containing 1% BME. T, S, and P samples were brought to 1x concentration of SDS sample buffer containing 1% BME (final). All samples were heated at 100°C for 5 min followed by centrifugation at 10,000 x *g* for 1 min. 15 *μ*L of (UN), (IN) and 30 *μ*L of (T), (P), (S) were resolved in 10% denaturing gels and the gels were stained with GelCode Blue Stain Reagent (Thermo Fisher Scientific) and imaged.

### Aggregation assays of proteasome subunits in yeast

Yeast cultures were grown in YPD or synthetic defined media (with casamino acids) overnight. The next day, cultures were diluted to OD_600_=0.2 in YPD or synthetic defined media (with casamino acids) and grown to mid-log or saturation phase. Cells were harvested and washed with sterile cold water and flash frozen in liquid N_2_. Cells were ground using a mortar and pestle until a fine powder was formed. Cell powder was resuspended in ice-cold lysis buffer (50 mM Tris pH 7.5, 150 mM NaCl, 1% glycerol, 1 mM EDTA, 1 mM PMSF, 1X EDTA-free cOmplete Protease Inhibitor Cocktail (Roche)) and vortexed intermittently during a 10 min incubation on ice. Samples were centrifuged at 3000 x *g* for 30 s to remove unlysed cells and cell debris.

Supernatants were transferred to fresh tubes. BCA assays were conducted to determine total protein concentrations. Protein concentration was normalized across all samples tested. A small aliquot was set aside as total protein (T). The remaining normalized supernatants were centrifuged at 100,000 x *g* for 20 min at 4°C in a Beckman Coulter TLA-55 rotor. Supernatants (S) were transferred to fresh tubes. Pellets were then washed with lysis buffer and re-centrifuged as above. Supernatants were discarded and the resulting pellets were resuspended in half the volume of the supernatant (S) to make (2xP). (T), (S), and (2xP) samples were brought to 1x concentration of SDS sample buffer containing 1% BME (final). Samples were heated at 100°C for 5 min followed by centrifugation at 10,000 x *g* for 1 min. Equal volumes of T, S, and 2xP samples were loaded onto 10% SDS gels and subjected to immunoblot analyses.

### Antibodies and immunoblotting

After samples were resolved in denaturing SDS or non-denaturing polyacrylamide gels, proteins in the gels were transferred to PVDF membranes (Millipore). Immunoblots were analyzed using primary antibodies against α4/Pre6 (D. Wolf), Rpt1 (W. Tansey), Rpt2 (Enzo Life Sciences), Rpt3 (Enzo Life Sciences), Rpt4 (W. Tansey), Rpt5 (Enzo Life Sciences), Rpn2 (M. Glickman), Rpn12 (D. Finley), ubiquitin (Dako), phosphoglycerate kinase (PGK; Invitrogen), and glucose-6-phosphate dehydrogenase (G-6-PDH; Sigma-Aldrich). For enhanced chemiluminescence detection (ECL), horseradish peroxidase (HRP)-linked anti-mouse IgG (from sheep) and HRP-linked anti-rabbit IgG (from donkey) (both GE Healthcare) were used as secondary antibodies.

### Analyses of mRNA levels in yeast extracts

Yeast cultures were grown in selective defined media overnight. Cultures were diluted to OD_600_=0.2 in selective defined media and grown to mid-log phase. Cells corresponding to one OD_600_ unit were harvested and washed with sterile ice-cold water. Total RNA was extracted from the cells using an RNeasy Mini Kit (Qiagen) and eluted in 50 *μ*L nuclease-free water. Contaminating DNA was subsequently removed from the samples using the DNA-*free*^TM^ Kit (Ambion). Two *μ*g of total RNA was reverse transcribed using the iScript^TM^ cDNA Synthesis Kit (Bio-Rad). The resulting cDNA was subjected to quantitative PCR reactions using iQ SYBR Green Supermix (Bio-Rad) and analyzed on a LightCycler 480 (Roche). Each qPCR reaction was conducted in three technical triplicates. All experiments were conducted according to manufacturers’ instructions.

### Ubiquitin pulldown assays

To determine if proteasome subunits were ubiquitinated at non-permissive temperature, we conducted a ubiquitin pulldown assay with slight modifications from a previously outlined protocol (40). We transformed WT and *rpt2,5PA* strains with pUB175 (expressing untagged ubiquitin) or pUB221 (expressing His_6_-tagged ubiquitin). The yeast ubiquitin genes in these plasmids are expressed under the control of a copper-induced *CUP1* promoter. Overnight cultures grown at 30°C were diluted to OD=0.2 and grown in 175 mL SD-URA for 2.5 hours at 37°C. The cultures were then induced with 0.5 mM CuSO_4_ (final) and grown for another 4 hours. 5 mL of each culture were harvested, washed with sterile water, and set aside as input.

The remaining cultures were harvested by centrifugation, washed with sterile water, and resuspended in 2 mL Buffer A (6 M guanidine-HCl, 0.1 mM Na_2_HPO_4_/NaH_2_PO_4_, 10 mM imidazole, pH 8.0) followed by cell disruption with glass beads for 6 x 20 s at top speed with 30 s breaks on ice between each round. Samples were centrifuged at 1690 x *g* for 15 min. Supernatants were collected, and total protein concentration of each sample was determined via Bradford assay (Bio-Rad). Total protein was normalized to 2 mg across all samples tested and incubated with 0.25 mL of 50% Ni-NTA resin (Qiagen) for 2 hours. The resin was subsequently pelleted, and the supernatant was aspirated off. The remaining beads were washed three times with 1 mL Buffer A followed by three washes with 1mL Buffer A/TI (1 volume Buffer A and 3 volumes Buffer TI–25 mM Tris-HCl, 20 mM imidazole, pH 6.8) and finally once with 1mL Buffer TI. The beads were then resuspended in 0.20 mL 2x SDS sample buffer (containing 0.2 mM imidazole and 8% BME) and subsequently boiled for 5 min.

Input samples that had been set aside were lysed by resuspending the pellet in EZ buffer (0.06 M Tris-HCl pH 6.8, 10% glycerol, 2% SDS, 5% BME) and boiled for 10 min. Bradford assay was conducted to determine protein concentration, and 10 *μ*g of each sample were resuspended in 2x sample buffer (containing 0.2 mM imidazole and 8% BME) and further boiled for another five minutes. 10 *μ*g of input sample and 30 *μ*L of each pulldown sample were loaded onto 10% denaturing gels and subjected to immunoblot analyses.

## Data availability

All data are contained within this manuscript.

## Acknowledgements

We thank Carolyn Breckel, Hongli Chen, and Jianhui Li for critical reading of the manuscript. We are also grateful to Dan Finley, Michael Glickman, William Tansey, and Dieter Wolf for providing antibodies used in this study. This work was supported by National Institute of Health grants (GM083050 and GM136325) to M.H.

## Abbreviations

CC: coiled coil
OB: oligonucleotide-binding
PAN: proteasome-activating nucleotidase
CP: core particle
RP: regulatory particle
cryo-EM: cryogenic electron microscopy
PQC: protein quality control

## SUPPORTING INFORMATION

Figure S1: Genetic interactions between *rpt* P-to-A mutations and different base and CP assembly chaperone gene deletions

Figure S2: Genetic interactions between *rpn4*Δ and *rpt* P-to-A mutations

Figure S3. Growth analysis of *rpt* double P-to-A mutants

Figure S4. Growth analysis of *pre9Δ* and *sem1Δ* assembly mutants expressing *not4-L35A* ubiquitin ligase

Figure S5. Analysis of aggregation and ubiquitination of select proteasome base subunits in *rpt2,5PA*

Figure S6. Growth analysis of select base, CP, and lid assembly mutants with chromosomal *HSP42* gene deleted

Table S1. List of diverse eukaryotic species analyzed for phylogenetic analyses

Table S2. List of yeast strains used in this study

Table S3. List of plasmids used in this study

---

**Figure S1.**
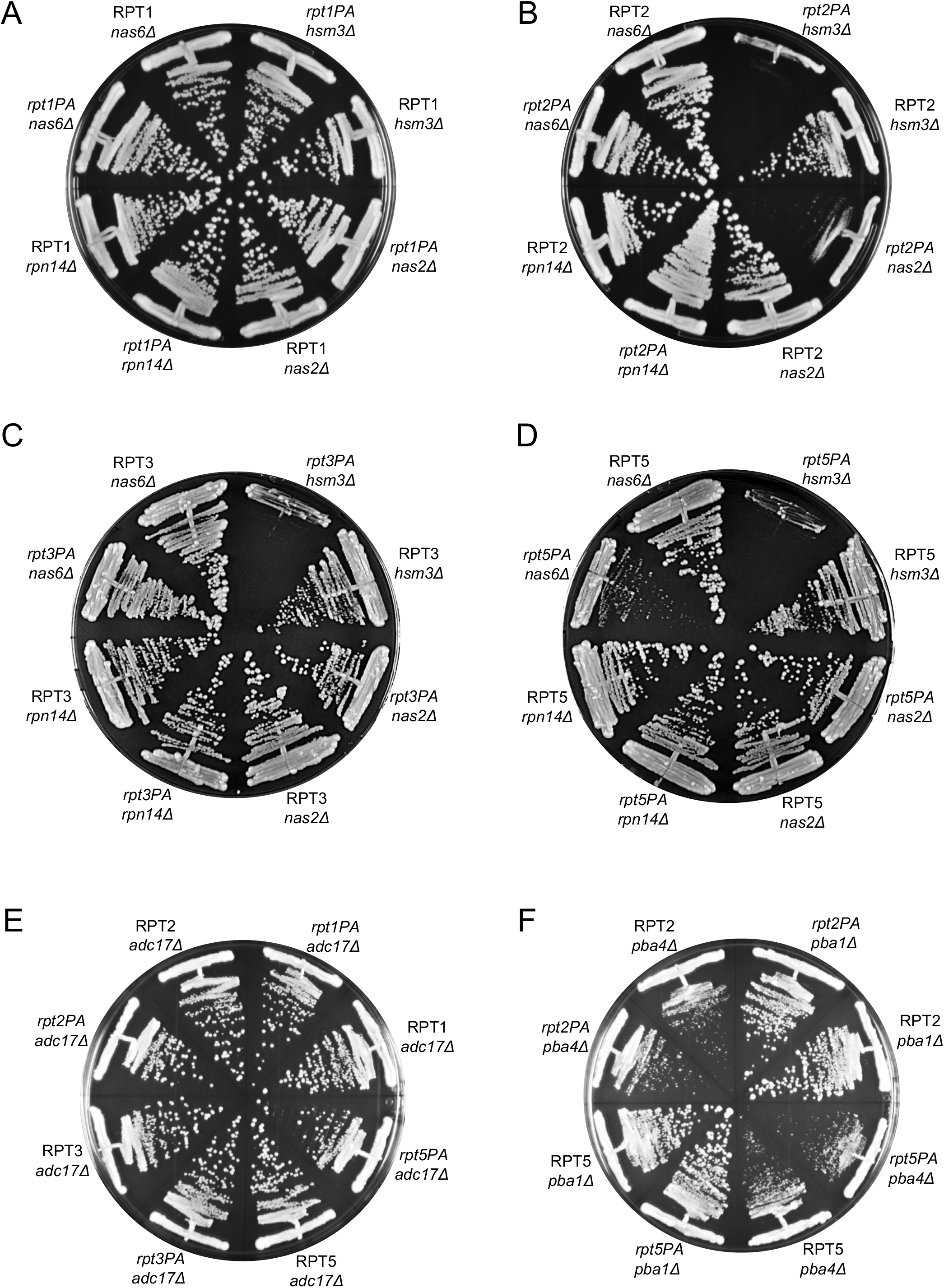
Growth assays evaluating potential synthetic interactions between single *rpt* P-to-A mutations and different base and CP assembly chaperone gene deletions. Cells were grown on YPD plates at 36°C.

**Figure S2.**
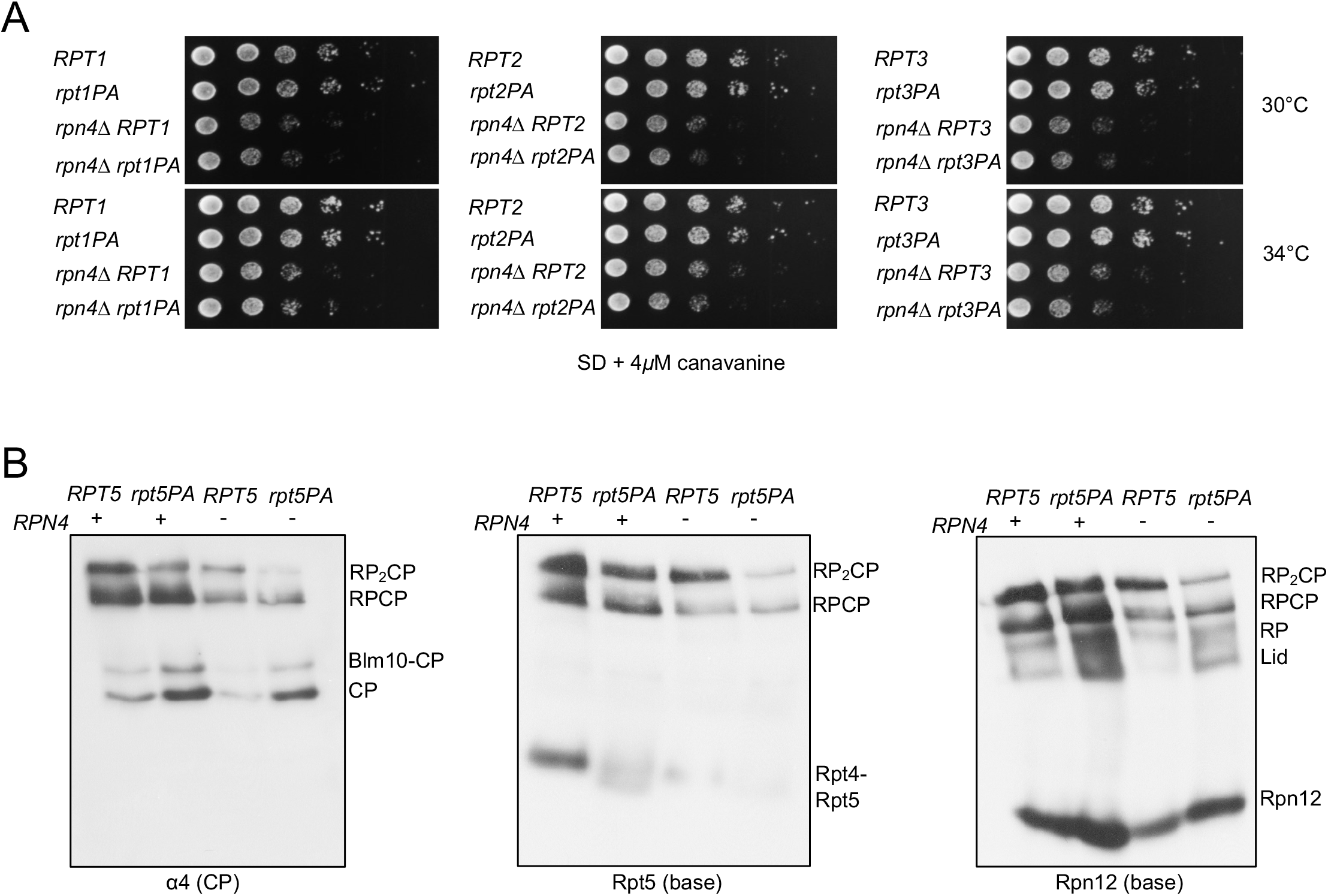
Genetic interactions between *rpn4*Δ and *rpt* P-to-A mutations. (A) No synthetic growth defects observed between *rpn4*Δ and *rpt1*-*P76A*, *rpt2-P103A*, or *rpt3-P96A.* Serially diluted cultures were spotted on plates as in Figure 2B. (B) Immunoblot analyses of yeast proteasome complexes in *rpt5-P76A* cells with and without deletion of *RPN4*.

**Figure S3.**
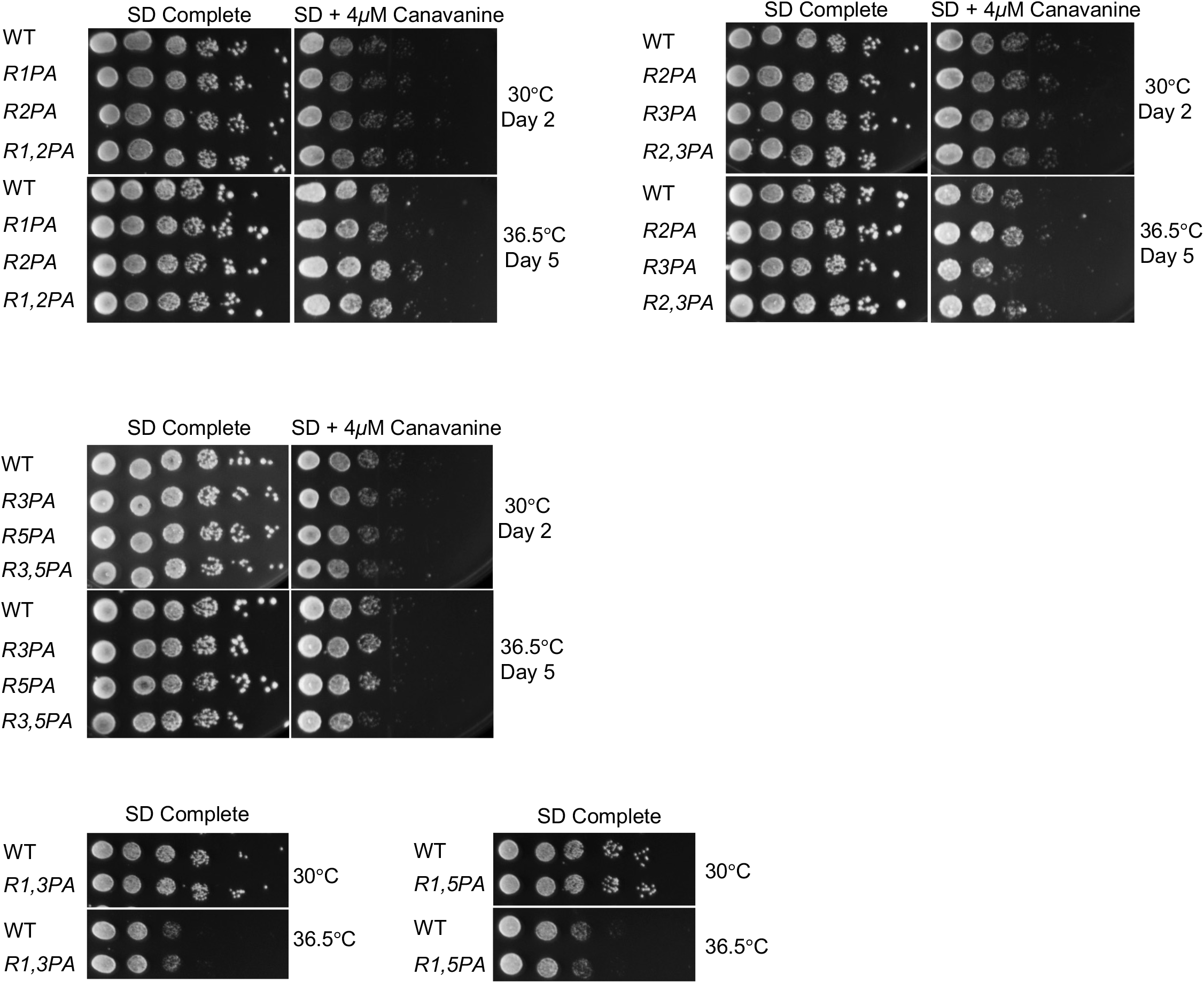
No significant growth defects are observed in the indicated *rpt* double P-to-A mutants. Serially diluted cultures were spotted on plates as in Figure 2B.

**Figure S4.**
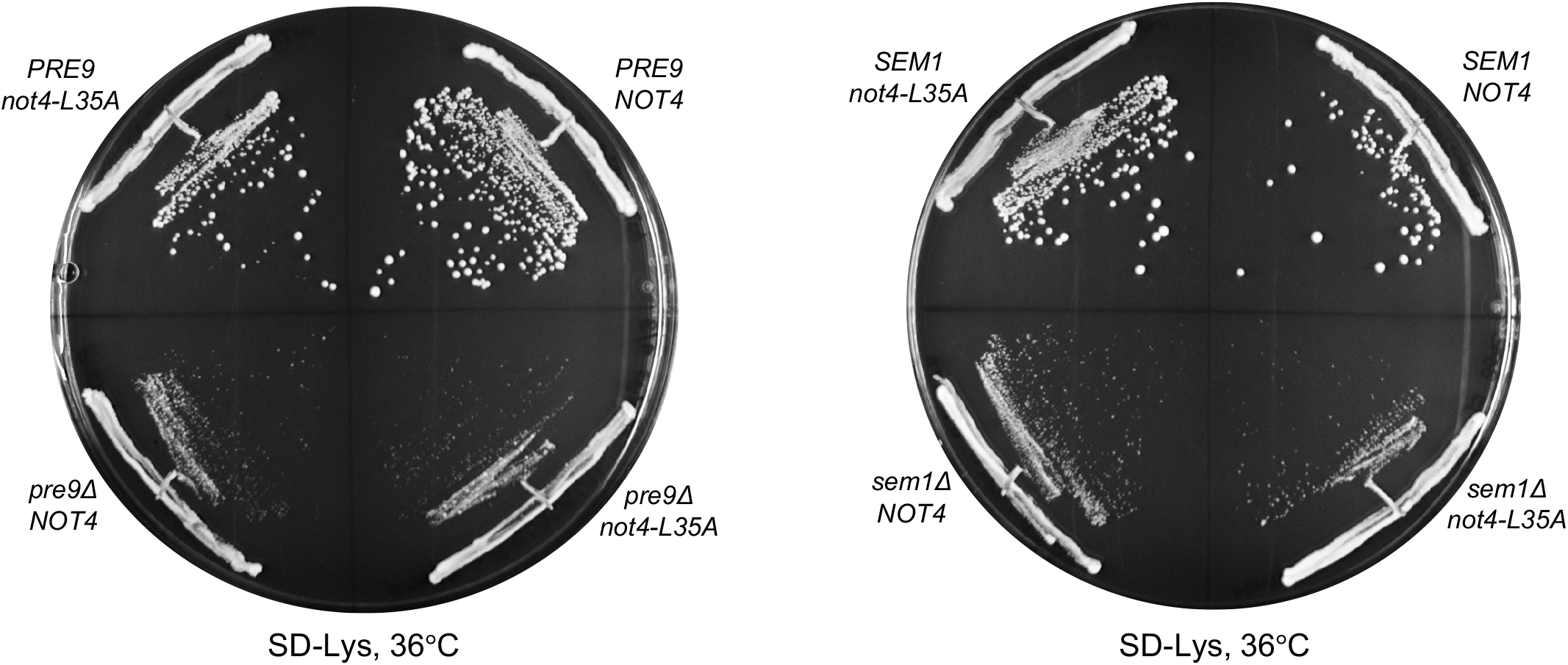
The *not4-L35A* ubiquitin ligase mutation does not rescue the tested CP (*pre9Δ*) and RP (*sem1Δ*) assembly mutants

**Figure S5.**
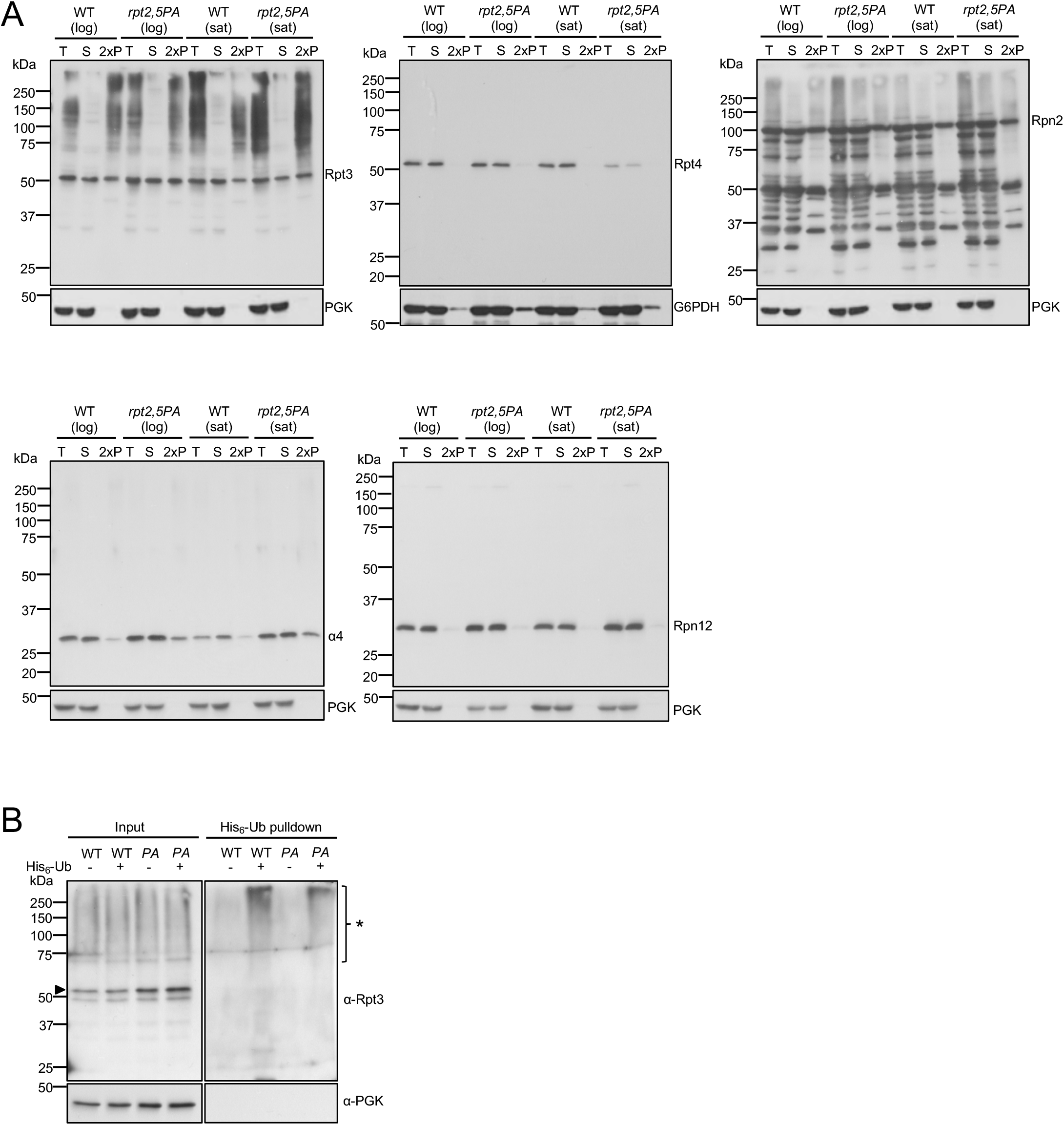
Analysis of aggregation and ubiquitination of select proteasome base subunits in *rpt2,5PA* mutant cells. (A) Yeast aggregation assays in *rpt2,5PA* versus WT cells. The data suggest that aggregation of the base subunit Rpt3 increases in the mutant but base subunit Rpn2 aggregation is unaffected. Bulk Rpt4 steady-state levels are notably decreased in *rpt2,5PA* when cells are grown to saturation, which is accompanied by decreased overall translation. CP subunit α4 aggregation is not obviously affected by *rpt2,5PA*, while aggregation of lid subunit Rpn12 is also unaffected. (B) Purification of bulk His6-ubiquitin conjugates reveals comparable Rpt3 ubiquitination in both WT and *rpt2,5PA* strains at 37°C as in Figure 5B. Arrowhead denotes unmodified Rpt3 bands. *, Ubiquitinated Rpt3 species.

**Figure S6.**
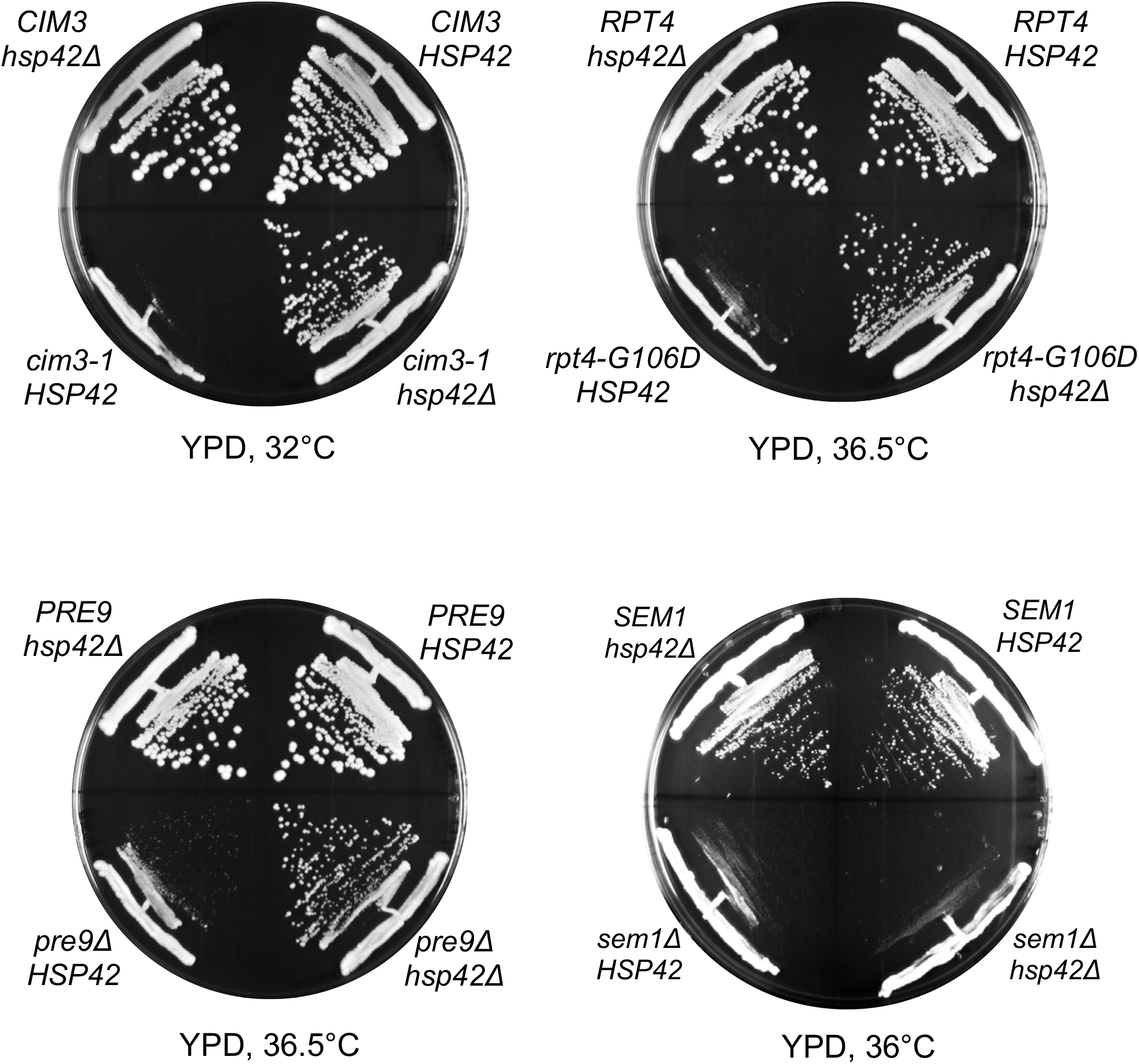
*HSP42* deletion suppresses growth defects of other base (*cim3-1* and *rpt4-G106D*) and CP (*pre9Δ*) assembly mutants but not that of a *sem1Δ* (lid) mutant

**Table S1.**
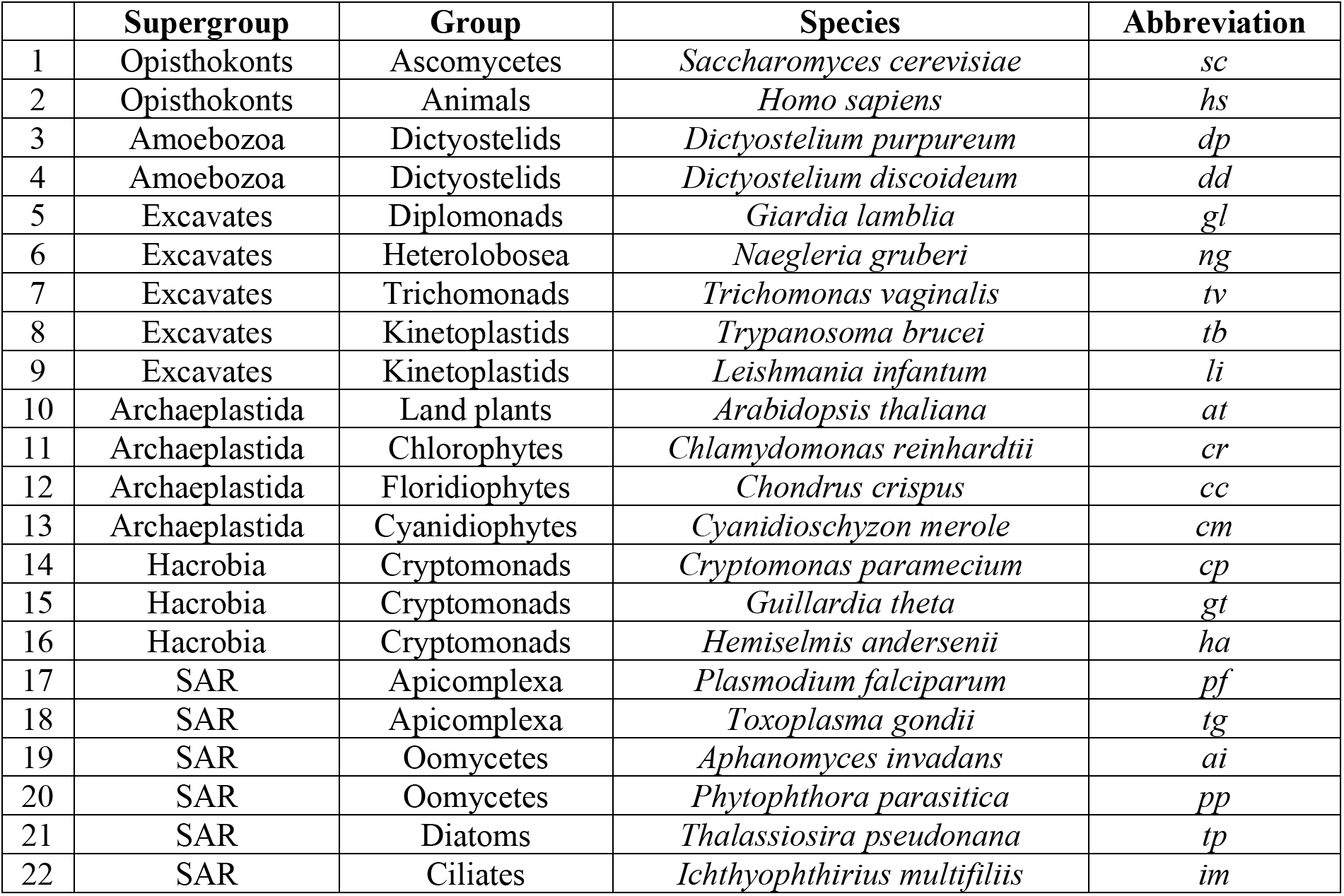
List of diverse eukaryotic species analyzed for phylogenetic analyses

**Table S2:**
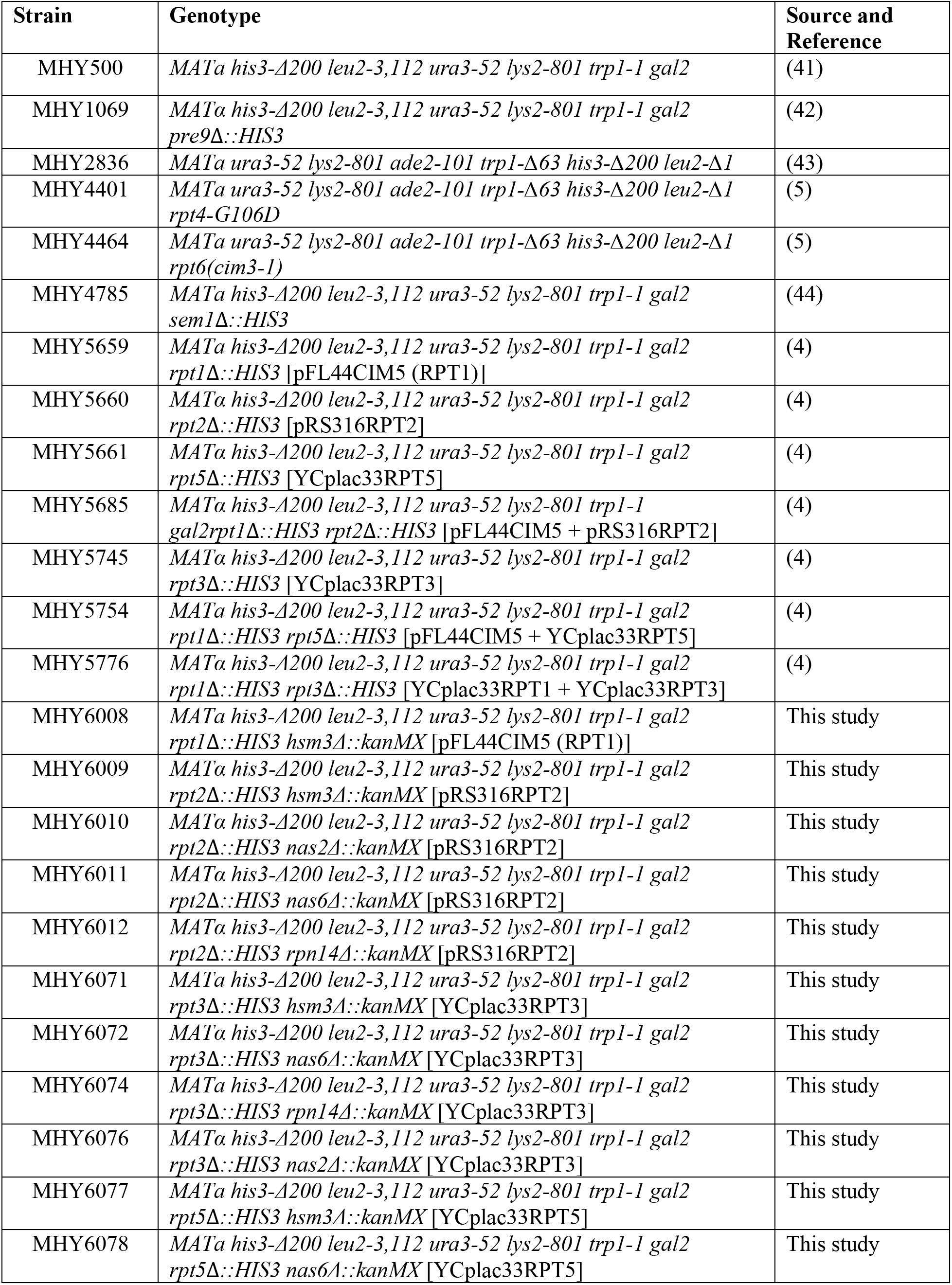
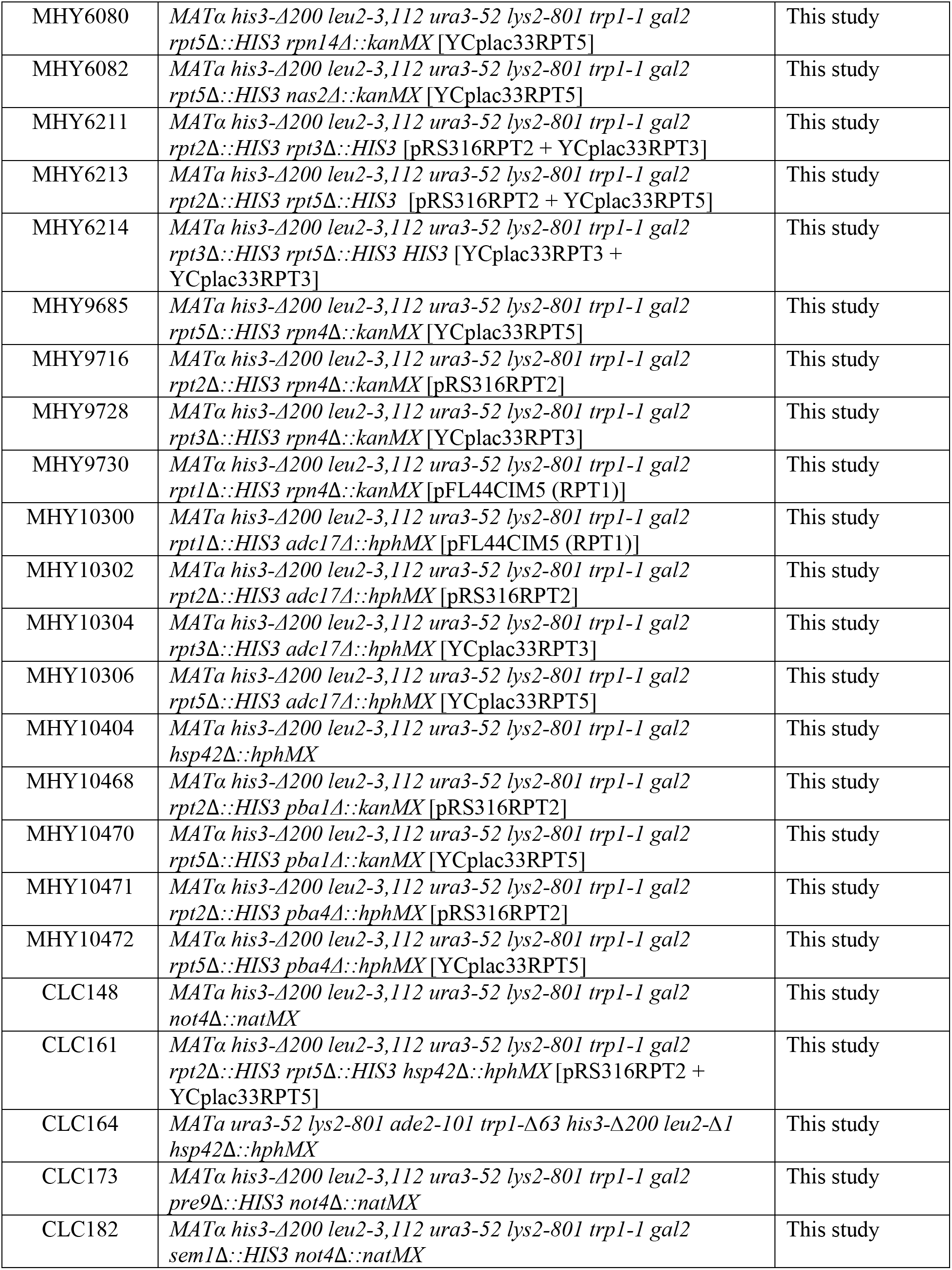
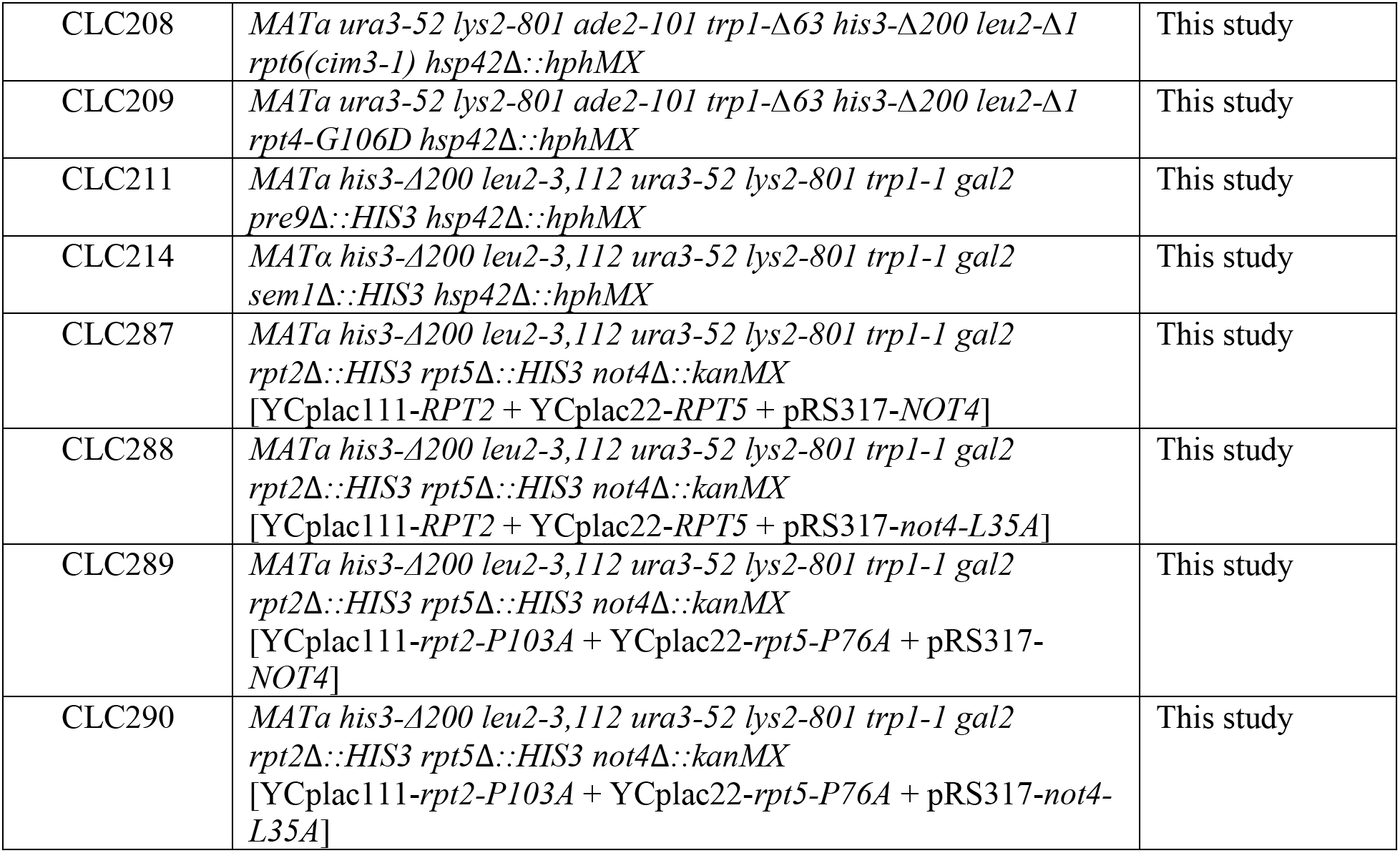
List of yeast strains used in this study

**Table S3:**
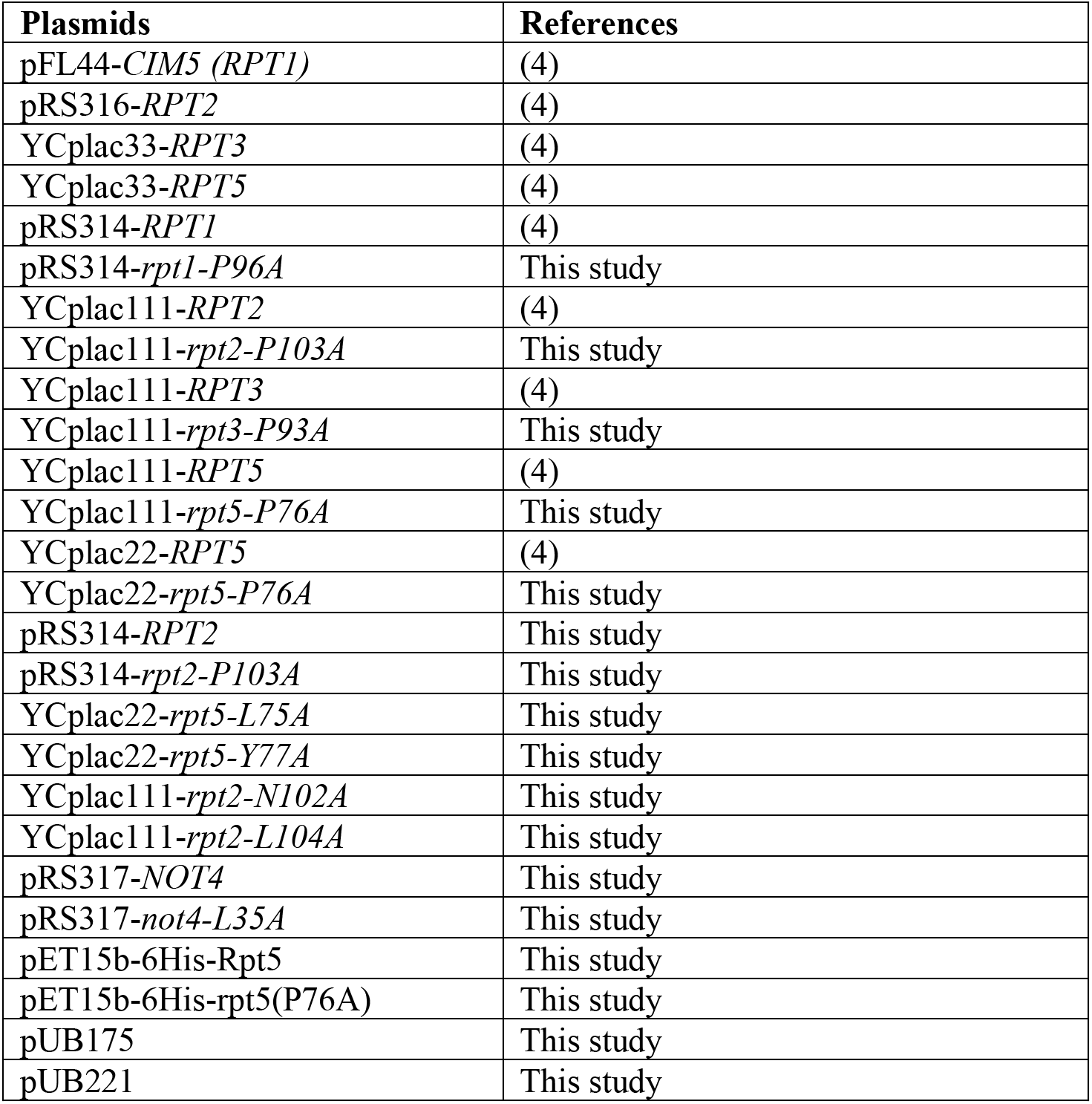
List of plasmids used in this study

| **Plasmids** | **References** |
| --- | --- |
| pFL44-*CIM5 (RPT1)* | (4) |
| pRS316-*RPT2* | (4) |
| YCplac33-*RPT3* | (4) |
| YCplac33-*RPT5* | (4) |
| pRS314-*RPT1* | (4) |
| pRS314-*rpt1-P96A* | This study |
| YCplac111-*RPT2* | (4) |
| YCplac111-*rpt2-P103A* | This study |
| YCplac111-*RPT3* | (4) |
| YCplac111-*rpt3-P93A* | This study |
| YCplac111-*RPT5* | (4) |
| YCplac111-*rpt5-P76A* | This study |
| YCplac22-*RPT5* | (4) |
| YCplac22-*rpt5-P76A* | This study |
| pRS314-*RPT2* | This study |
| pRS314-*rpt2-P103A* | This study |
| YCplac22-*rpt5-L75A* | This study |
| YCplac22-*rpt5-Y77A* | This study |
| YCplac111-*rpt2-N102A* | This study |
| YCplac111-*rpt2-L104A* | This study |
| pRS317-*NOT4* | This study |
| pRS317-*not4-L35A* | This study |
| pET15b-6His-Rpt5 | This study |
| pET15b-6His-rpt5(P76A) | This study |
| pUB175 | This study |
| pUB221 | This study |

